# An ‘invisible’ ubiquitin conformation is required for efficient phosphorylation by PINK1

**DOI:** 10.1101/189027

**Authors:** Christina Gladkova, Alexander F. Schubert, Jane L. Wagstaff, Jonathan N. Pruneda, Stefan M.V. Freund, David Komander

## Abstract

The Ser/Thr protein kinase PINK1 phosphorylates the well-folded, globular protein ubiquitin (Ub) at a relatively protected site, Ser65. We had previously shown that Ser65-phosphorylation results in a conformational change, in which Ub adopts a dynamic equilibrium between the known, common Ub conformation and a distinct, second conformation in which the last β-strand is retracted to extend the Ser65 loop and shorten the C-terminal tail. We here show using Chemical Exchange Saturation Transfer (CEST) NMR experiments, that a similar, C-terminally retracted (Ub-CR) conformation exists in wild-type Ub. Ub point mutations in the moving β5-strand and in neighbouring strands shift the Ub/Ub-CR equilibrium. This enabled functional studies of the two states, and we show that the Ub-CR conformation binds to the PINK1 kinase domain through its extended Ser65 loop and is a superior PINK1 substrate. Together our data suggest that PINK1 utilises a lowly populated yet more suitable Ub-CR conformation of Ub for efficient phosphorylation. Our findings could be relevant for many kinases that phosphorylate residues in folded proteins or domains.

## INTRODUCTION

Protein ubiquitination and protein phosphorylation are the two main regulatory post-translational modifications of proteins (Hunter, 2007). While phosphorylation provides a binary signal, the ubiquitin (Ub) signal is highly tuneable and exists in many variations. For example, polyUb chains of many architectures exist and encode distinct biological outcomes (Komander & Rape, 2012); moreover, Ub itself can be phosphorylated or acetylated, expanding its functional versatility (Swatek & Komander, 2016; Yau & Rape, 2016). Mass-spectrometry has enabled the discovery and quantitation of the plethora of Ub modification sites (Ordureau *et al*, 2015), yet proteins regulating and responding to these have remained by-and-large unclear. So far, the best studied Ub modification is Ser65-phosphorylated Ub (hereafter phosphoUb), which has been intimately linked to mitophagy, the process by which mitochondria isolate damaged parts and target them for autophagic clearance (Nguyen *et al*, 2016; Pickrell & Youle, 2015).

PhosphoUb is generated on mitochondria by the Ser/Thr protein kinase PINK1 (Kazlauskaite *et al*, 2014; Koyano *et al*, 2014; Kane *et al*, 2014; Ordureau *et al*, 2014; Wauer *et al*, 2015b), which is stabilised on the cytosolic face of mitochondria upon membrane depolarisation (Narendra *et al*, 2010). PINK1 phosphorylates Ub attached to outer mitochondrial membrane proteins, and this recruits and allosterically activates the E3 ligase Parkin (Wauer *et al*, 2015a; Kazlauskaite *et al*, 2015; Sauvé *et al*, 2015; Kumar *et al*, 2015). PINK1 also phosphorylates Parkin in its Ub-like (Ubl) domain, which is required for full Parkin activation and leads to strong, localised mitochondrial ubiquitination (Kondapalli *et al*, 2012; Wauer *et al*, 2015a; Ordureau *et al*, 2014). PINK1/Parkin action attracts adaptor proteins and recruits the mitophagy machinery, leading to clearance of the damaged organelle (Lazarou *et al*, 2015; Heo *et al*, 2015). The pathophysiological importance of PINK1/Parkin mediated mitophagy is underlined by the fact that mutations in PINK1 and Parkin are linked to autosomal recessive juvenile Parkinson’s disease (AR-JP), a neurodegenerative condition arising from loss of dopaminergic neurons in the substantia nigra (Corti *et al*, 2011; Pickrell & Youle, 2015).

The generation of phosphoUb by PINK1 is mechanistically poorly understood. PINK1 is an unusual Ser/Thr kinase, highly divergent from other kinases in the kinome (Manning *et al*, 2002). In part this is due to several large insertions in the kinase N-lobe, which complicate structural modelling (Trempe & Fon, 2013). Also its substrate, Ub, is a non-classical kinase target since its 76 amino acids form a globular, highly robust and stable β-grasp fold, in which Ser65 is markedly protected. Ub Ser65 resides in the loop preceding the β5-strand, and its side chain hydroxyl group engages in a backbone hydrogen bond with Gln62. In addition, nearby side chains of Phe4 and Phe45 further stabilise the Ser65-containing loop (**Fig. EV1A**). Ub Ser65 is structurally identical to Ser65 in the Parkin Ubl domain, but the two substrates lack similarity at the sequence level and a PINK1 phosphorylation consensus motif is not obvious (Kazlauskaite *et al*, 2014). The Ser65 position and interactions within a well-folded, globular domain make this residue an unlikely phosphorylation site for PINK1 or indeed any kinase.

Ub is highly similar in the >100 Ub crystal structures in the protein data bank (Perica & Chothia, 2010), and its biophysical properties and availability have made it a popular model system for protein folding studies (Jackson, 2006) and nuclear magnetic resonance (NMR) techniques (Torchia, 2015; Lange *et al*, 2008; Fushman *et al*, 2004). NMR studies in particular have shown that despite its compact fold and high intrinsic stability, Ub is very dynamic and contains several regions of local conformational flexibility (Lange *et al*, 2008). These include a mobile four-residue C-terminal tail, as well as a flexible β hairpin structure, the β 1/ β 2-loop, that alters the interaction profile of Ub (Lange *et al*, 2008; Hospenthal *et al*, 2013; Phillips & Corn, 2015). Importantly, we previously discovered that Ser65-phosphorylation resulted in a further, dramatic conformational change in Ub (Wauer *et al*, 2015b).

The Ser65 loop and the last β5 strand were previously not known to be conformationally dynamic, yet phosphorylation led to an equilibrium between two phosphoUb conformations (**Fig. 1A**). The first state resembles the ‘common’ Ub conformation observed all reported crystal structures to date (hereafter simply Ub conformation). This phosphoUb conformation was confirmed in a crystal structure (Wauer *et al*, 2015b) and more recently by an NMR structure (Dong *et al*, 2017). More striking was a second conformation, in which the entire last β-strand slipped by two amino acids, extending the Ser65 loop, and simultaneously shortening the Ub C-terminal tail (hereafter referred to as the Ub-CR conformation for C-terminally retracted) (Wauer *et al*, 2015b; Dong *et al*, 2017). This change is facilitated by a Leu-repeat pattern in the β5-strand: Leu67, Leu69 and Leu71 occupy complementary Leu-pockets in the Ub core, whereas Leu73 is mostly solvent exposed. In the Ub-CR conformation observed in phosphoUb, Leu73 occupies the Leu71 pocket, Leu71 occupies the Leu69 pocket, and Leu69 occupies the Leu67 pocket, resulting in Leu67 residing in a more exposed position that was formerly occupied by Ser65 (**Fig. 1A**). Experimentally, the phosphoUb-CR conformation was supported by large (>1.5 ppm) chemical shift perturbations and by determination of the hydrogen bonding patterns for the β-sheet, using long range HNCO-based NMR analysis (Wauer *et al*, 2015b). A recent NMR structure of the phosphoUb-CR conformation confirmed our findings (Dong *et al*, 2017). We speculated that the Ub-CR conformation is a consequence of Ub phosphorylation, enabling proteins to specifically recognize phosphoUb (Wauer *et al*, 2015b).

**Figure 1.**
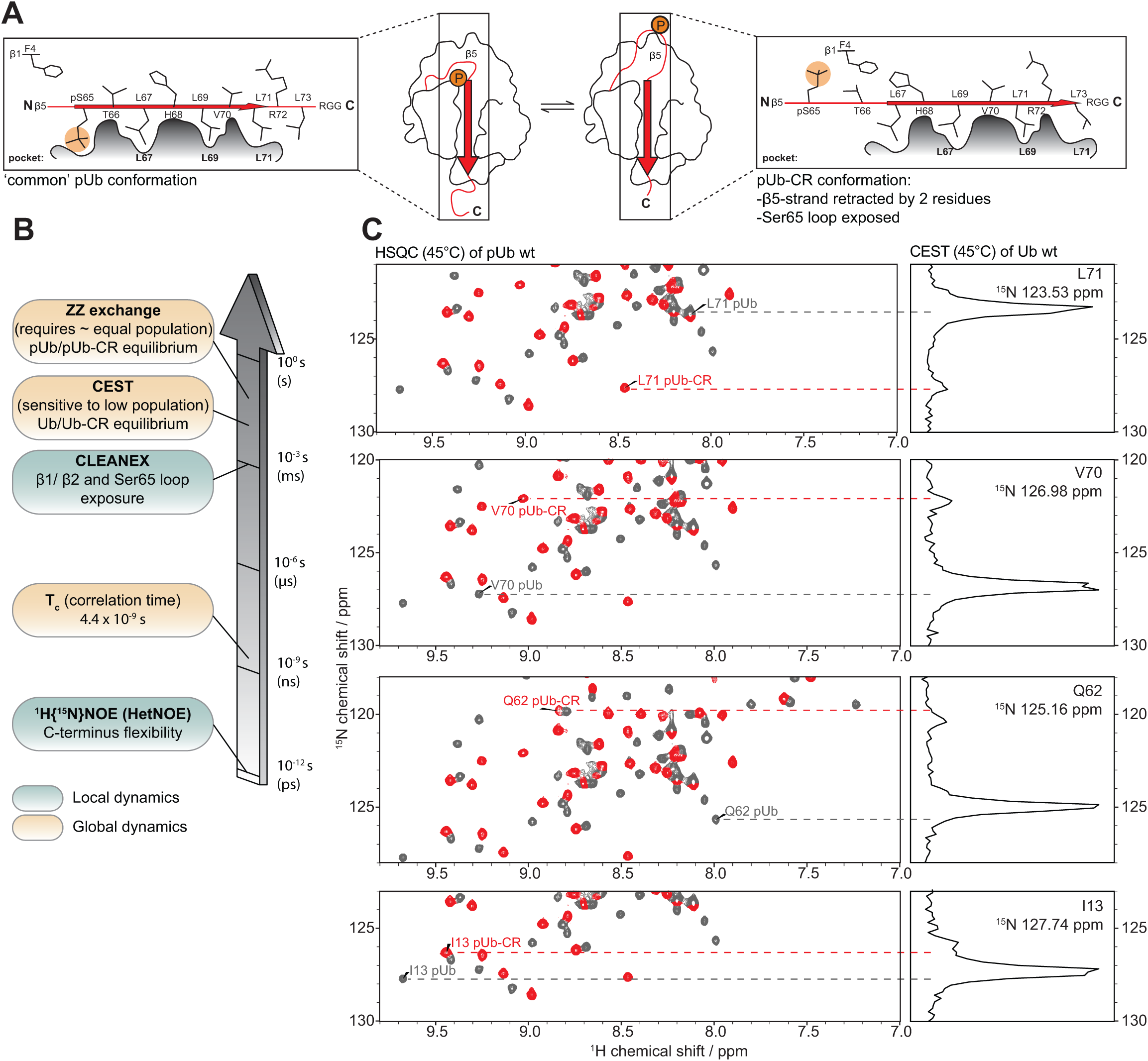
phosphoUb adopting the Ub C-terminally retracted (Ub-CR) conformation. ***A)*** *Centre*, Schematic of the Ub surface, showing the position of the β5-strand (arrow) on the Ub core, and the position of phosphorylated Ser65 is indicated. Cartoons to the left and right show a slice along the β-5 strand, depicting theβ-5 residues and their positions in the respective Ub conformation Leu pockets. **B)** Timescales of NMR experiments to study the Ub/Ub-CR conformation in this and previous work (Wauer *et al*, 2015b). **C)** CEST experiment on ^15^N-labelled wt Ub (1.5 mM) in phosphate buffered saline (pH 7.2) at 45 °C. For a subset of resonances in the HSQC spectrum of Ub, a cross section taken at their ^15^N frequency will display an additional resonance in this frequency-swept 2^nd^ ^15^N dimension (CEST profile) corresponding to the lowly populated Ub-CR conformation. The main peak in the CEST profile closely correlates to the corresponding HSQC resonance in the phosphoUb conformation (grey), while the amplified smaller peak matches the resonance position of the phosphoUb-CR conformation (red). Note that the observed chemical shift positions in the wt-Ub CEST data do not perfectly match phosphoUb resonances due to the chemical shift contribution of the phosphate group. Additional peaks can be found in **Fig. EV1B**, and full spectra in **Appendix Fig. S1**. A temperature profile for a selected resonance as well as a plot of all absolute ^15^N shift differences can be found in **Fig. EV1C and D**.

We here show that the Ub-CR conformation can indeed be detected in unphosphorylated Ub, when analysing NMR exchange timescales accessible by Chemical Exchange Saturation Transfer (CEST) experiments (**Fig. 1B**). This previously unrecognised equilibrium between a common Ub and a Ub-CR conformation in wild-type (wt) Ub can be shifted in either direction through point mutations in unphosphorylated Ub. Crystal structures as well as biophysical and NMR measurements enable in-depth characterisation of the Ub-CR conformation, and reveal its functional relevance during Ser65 phosphorylation. The Ub-CR conformation of Ub, with its mobile Ser65 loop, forms a more stable complex with PINK1 as assessed by NMR binding studies. More importantly, Ub in the Ub-CR conformation is a superior substrate for PINK1, and we provide evidence that the Ub-CR conformation is required for efficient PINK1 phosphorylation. Together, our work suggests that PINK1 utilises a lowly populated, ‘invisible’ form of Ub in which the buried phosphorylation site becomes exposed.

## RESULTS

### Identification of a Ub-CR conformation in wild-type Ub

We had previously shown that phosphoUb resides in a conformational equilibrium in which the β5-strand (aa 67-71) slides in its binding groove (**Fig. 1A**). The immediate question arising, even during the review of our earlier paper (Wauer *et al*, 2015b), was whether a similar Ub-CR conformation may exist in unphosphorylated, wt Ub.Dynamic aspects of Ub have been under excessive scrutiny, in particular by NMR, and numerous studies have collectively covered most motional timescales from fast ps internal motions up to μs-ms conformational exchange processes using RDC analysis (Lange *et al*, 2008; Torchia, 2015). Our initial detection of the phosphoUb / phosphoUb-CR transition was enabled by a near-equal population of both states, and ZZ-exchange experiments indicated a slow exchange (2 s^-1^) between these conformations (**Fig. 1B**).

Given the timescales of motion probed in previous Ub studies, we hypothesised that a very lowly populated, slow exchanging Ub-CR conformation in wt Ub could have been systematically missed in previous experiments. Furthermore, we assumed that raising the temperature of the experiment would lower the energy barrier between the two states and potentially allow an increase in population of the Ub-CR species. The detection of lowly populated, also known as ‘dark’ or ‘invisible’, conformational states can be enabled by CEST experiments (Vallurupalli *et al*, 2012; Kay, 2016). In CEST, samples are pulsed at specific frequencies to saturate individual resonances in the ‘invisible’ state. During an exchange period, magnetisation is transferred between conformers resulting in an amplification of the signals from previously ‘invisible’ resonances of the lowly populated conformations. Indeed, ^15^N-CEST experiments optimised for slow exchange processes (see **Methods**), revealed the existence of a second set of peaks in the ^15^N frequency sweep of wt Ub, under physiological conditions (phosphate buffered saline, pH 7.2 at 37 or 45 °C) (**Fig. 1C and EV1B-C**). The chemical shift positions of this second, lowly occupied population correlated well with previously recorded phosphoUb-CR resonances (**Fig. 1C, Fig. EV1B and D, Appendix Fig. S1**)

Together, CEST experiments revealed the existence of a previously undetected Ub conformation in wt Ub, which by chemical shift analysis resembles the phosphoUb-CR conformation reported earlier.

### Stabilisation of the Ub-CR conformation

With the occurrence of the wt Ub-CR conformation confirmed, we set out to stabilise it for further study. For this, we mutated Leu67 to Ser, hoping that occupation of the Leu67 pocket by Ser encourages β5-strand slippage to fill this hydrophobic pocket with Leu69 instead (**Fig. 2A**). Indeed, ^1H-15^N BEST-TROSY 2D spectra (bTROSY) of ^15^N-labelled Ub L67S showed 73 peaks implying a single Ub conformation (**Appendix Fig. S2**). The chemical shift pattern did not match wt Ub, but more closely resembled the pattern seen for the phosphoUb-CR conformation. This can be assessed using well-dispersed reporter resonances, such as Lys11 (**Fig. 2B**), while a global comparison of the full spectra can be drawn from chemical shift perturbation heat maps (**Fig. 2C**). Hence, Ub L67S predominantly adopts the Ub-CR conformation despite lacking phosphorylated Ser65.

**Figure 2.**
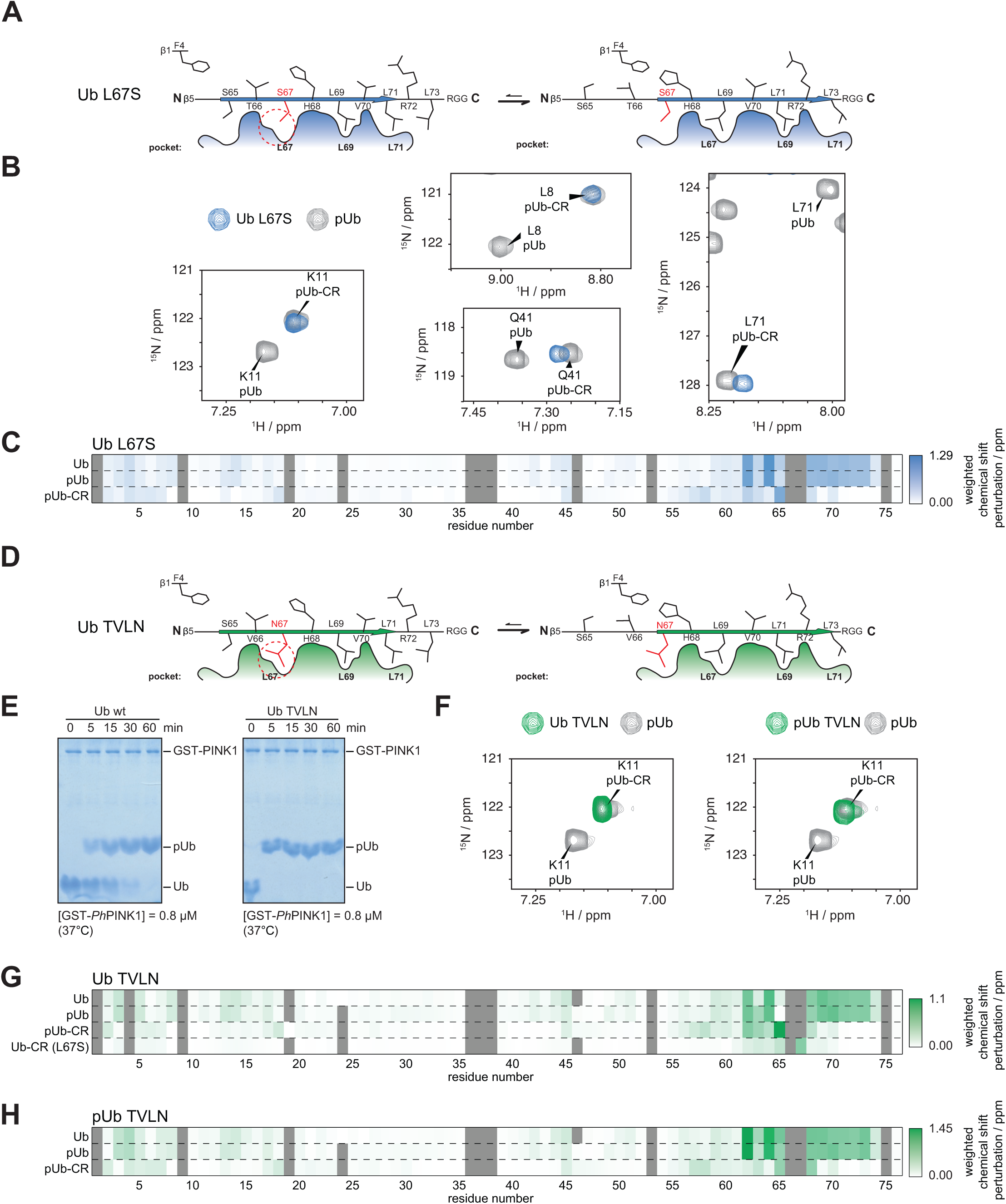
Stabilising the Ub-CR conformation with point mutations. **A)** Schematic of L67S mutation, which places a Ser in the Leu67 pocket. **B)** Selected resonances of Ub L67S, compared to previous phosphoUb / phosphoUb-CR spectra. Ub L67S adopts the Ub-CR conformation. For full spectra see **Appendix Fig. S2**. **C)** Weighted chemical shift perturbation heat maps, comparing Ub L67S to indicated Ub spectra, revealing the similarity with phosphoUb-CR.Schematic of Ub TVLN, introducing non-phosphorylatable residues at Thr66 and Leu67. For full spectra see **Appendix Fig. S3**. **D)** Phos-tag analysis of Ub TVLN phosphorylation by *Ph*PINK1. Like Ub, Ub TVLN is phosphorylated only on Ser65. **E)** Lys11 resonance of Ub TVLN in non- and phosphorylated state. Ub TVLN adopts only the Ub-CR conformation regardless of phosphorylation status. For full spectra see **Appendix Fig. S3**. **F)** Weighted chemical shift perturbation heat maps of Ub TVLN in comparison to indicated Ub species. **G)** Weighted chemical shift perturbation heat maps of phosphoUb TVLN in comparison to indicated Ub/phosphoUb species.

### Mimicking the Ub-CR conformer in Ser65 phosphoUb

We also wanted to study the phosphoUb-CR conformer in more detail, and hence we phosphorylated Ub L67S with *Pediculus humanus* (*Ph)*PINK1 (Wauer *et al*, 2015b; Woodroof *et al*, 2011). Strikingly, phosphorylation transformed the simple Ub-CR bTROSY spectrum to a complicated spectrum with the occurrence of many additional peaks (data not shown). Phos-tag gels and mass-spectrometry (MS) showed that *Ph*PINK1 phosphorylates Ub L67S at multiple sites, on Ser65, and on Thr66 or on the introduced Ser67 (**Fig. EV2A and B**). A mixture of phosphorylated species explains the complexity of the observed NMR spectrum. PINK1 shows exquisite preference for Ser65 in wt Ub, and only phosphorylates Thr66 at very high enzyme concentrations and late time points (Wauer *et al*, 2015b). Hence, the doubly phosphorylated species are a result of the Ub-CR conformation induced by the L67S mutation. These data indicated that the Ub-CR conformation has profound effects on PINK1-mediated Ub phosphorylation, but suggested that this mutation was limited in its usefulness for the study of phosphoUb-CR.

To overcome this and to generate exclusively Ser65-phosphorylated Ub in the Ub-CR conformation, Leu67 was mutated to Asn, and Thr66 was mutated to Val (termed hereafter Ub TVLN mutant) (**Fig. 2D**). Ub TVLN was phosphorylated only once, on Ser65 (**Fig. 2E**), showed a clean, single-species bTROSY spectrum highly similar to Ub L67S in the unphosphorylated form, and when phosphorylated was highly similar to the phosphoUb-CR conformation (**Fig. 2F-H**, **Appendix Fig. S3A and B**) (Wauer *et al*, 2015b). Together, this showed that phosphoUb TVLN is an excellent mimic for the Ub-CR conformer of the PINK1-catalysed phosphoUb species.

### Crystal structures of Ub in the Ub-CR conformation

The identification of Ub mutants stably in the Ub-CR conformation allowed us to obtain high-resolution crystal structures of Ub L67S (1.63 Å) and phosphoUb TVLN (1.6 Å) (**Table 1, Fig. 3, Fig. EV3**). Both structures confirmed that the β5-strand is retracted by two amino acids, and Ser/Asn67, Leu69, Leu71 and Leu73 adopt near identical conformations as compared to Ser65, Leu67, Leu69 and Leu71, respectively, seen in previous Ub structures (**Fig. 3A-D**). Hydrogen bonding patterns observed in the crystal structures matched the experimentally determined hydrogen bonding pattern for phosphoUb-CR conformation (Wauer *et al*, 2015b), and the phosphoUb TVLN structure is similar to a recently reported NMR structure of phosphoUb-CR (**Fig. EV3E and F**).

**TABLE 1.**
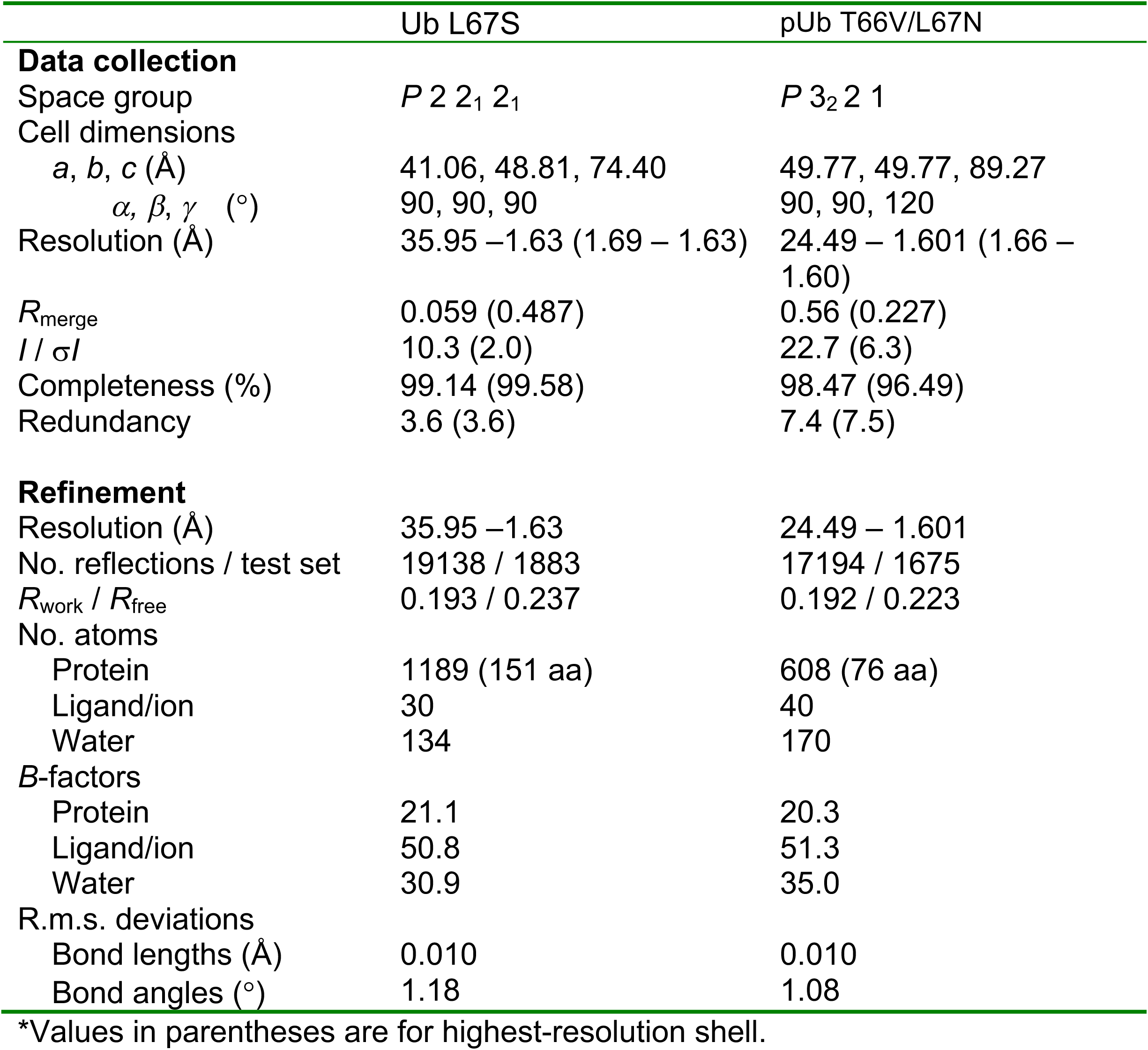
Data collection and refinement statistics.

**Figure 3.**
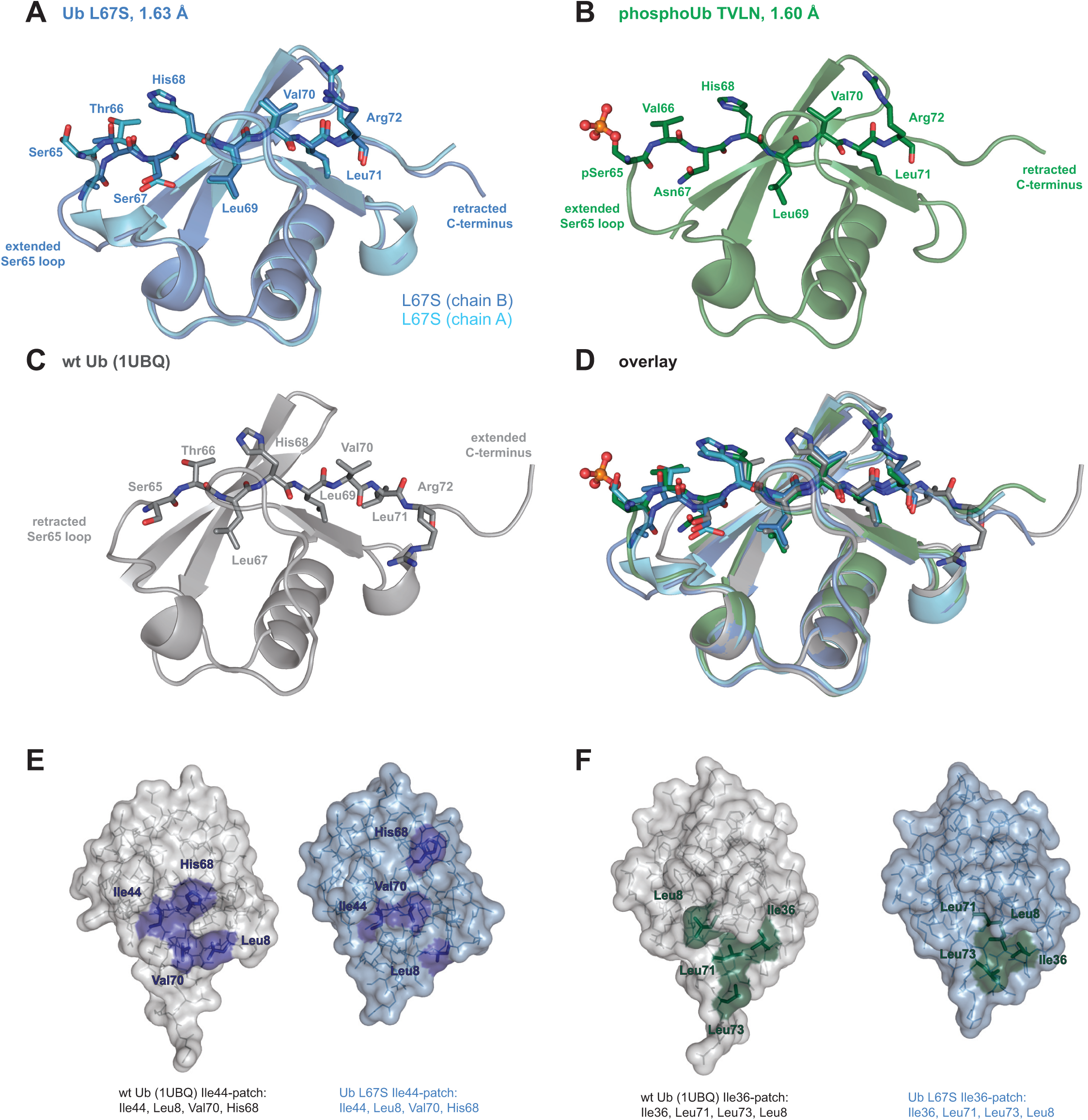
Crystal structures of Ub in the Ub-CR conformation. **A)** Ub L67S structure at 1.63 Å resolution. The two molecules of the asymmetric unit are superimposed. For electron density **see Fig. EV3A**. **B)** PhosphoUb TVLN structure at 1.6 Å resolution. For electron density **see Fig. EV3B**. **C)** Structure of wt Ub (PDB-id 1UBQ, (Vijay-Kumar *et al*, 1987)). **D)** Superposition of structures from A-C, showing residues of the β5-strand. **E)** Position of residues making up the Ile44 hydrophobic patch in Ub or Ub-CR conformations. **F)** As in E, showing residues of the Ile36 hydrophobic patch.

The structures reveal important consequences of the Ub-CR conformation. The Ser65-containing loop (aa 62-66) protrudes from and lacks defined contacts with the Ub core, is flexible judging by B-factor analysis, and in Ub L67S adopts distinct conformations in the two molecules in the asymmetric unit (**Fig. 3A, Fig. EV3D**). Likewise, in phosphoUb TVLN, the Ser65-containing loop is extended and seemingly mobile, and the phosphate group is exposed and does not contact the Ub core. A further important feature of the Ub-CR conformation is the disruption of Ub interaction interfaces, the most important of which is the Ile44 hydrophobic patch, which also utilizes Leu8 in the flexible β1/β2-hairpin, and Val70 and His68 of Ub β5-strand (Komander & Rape, 2012). In the Ub-CR conformation, the Ile44 hydrophobic patch is disrupted due to dislocation of β5-residues Val70 and His68 (**Fig. 3E**). In contrast, a second interaction site, the Ile36 hydrophobic patch (Hospenthal *et al*, 2013) is intact (**Fig. 3F, Fig. EV3C**). Finally, retraction of β5-strand by two residues reduces the reach and conformational flexibility of the important Ub C-terminal tail.

### Affecting Ub/Ub-CR conformational equilibrium

Mutating the first hydrophobic residue of the β5 strand, Leu67, favours the Ub-CR conformation, since Leu69 and Leu71 can occupy alternative positions easily. We reasoned that mutating Leu71 to a larger residue, which cannot occupy the Leu69 position, might stabilise it in the common Ub conformation, and disfavour the Ub-CR conformation after phosphorylation (**Fig. 4A**). Indeed, this was the case; Ub L71Y displays a common Ub spectrum without phosphorylation, and a spectrum highly similar to the common phosphoUb species after phosphorylation (**Fig. 4B-D, Appendix Fig. S4A and B)**. Hence, Ub L71Y is a mutation in which the Ub-CR conformation is disfavoured.

**Figure 4.**
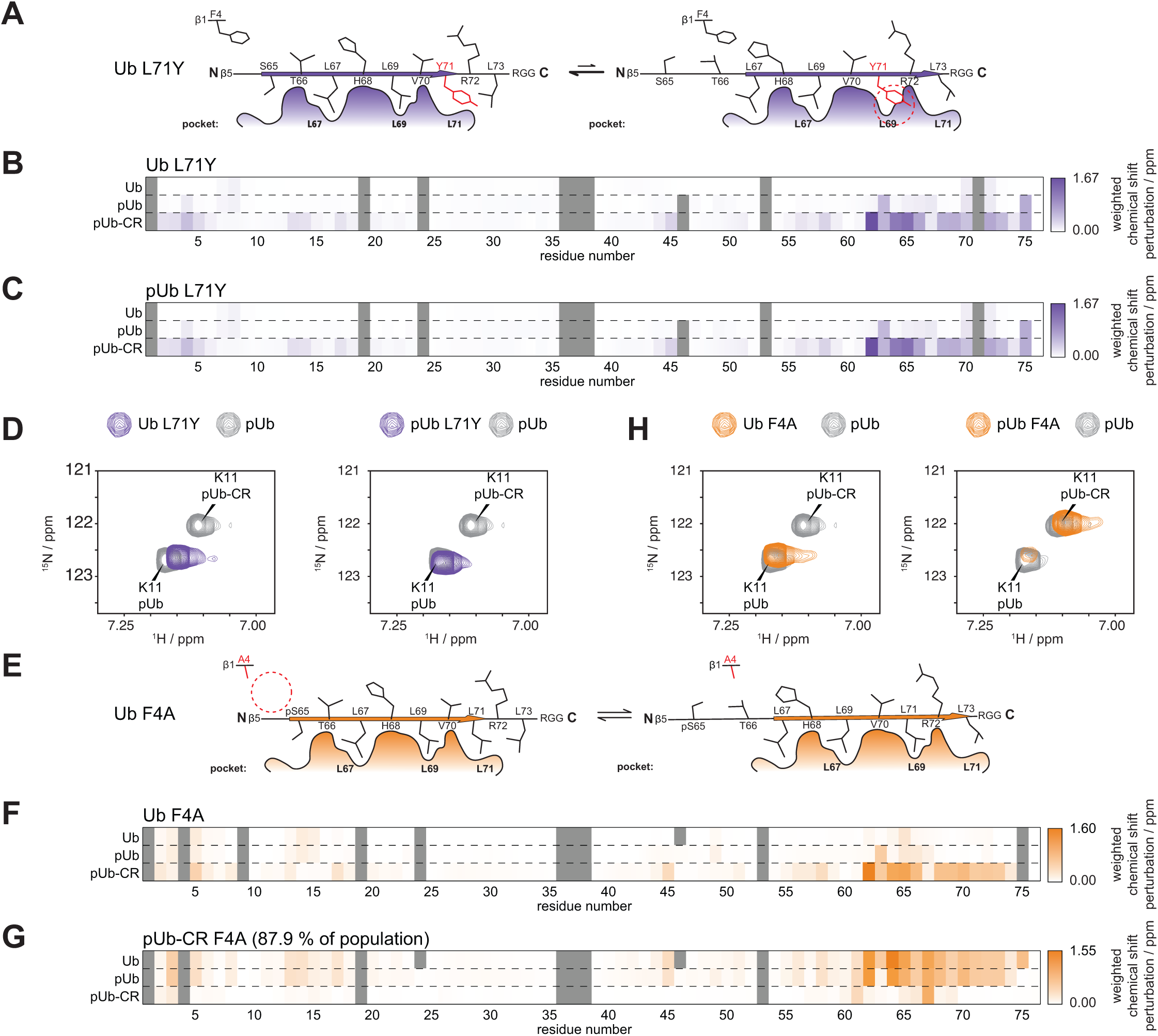
Mutations to modulate the Ub / Ub-CR equilibrium. **A)** Schematic of the Ub L71Y mutation. A large Tyr residue may not easily fit into the Leu69 pocket. **B)** Weighted chemical shift perturbation heat maps of Ub L71Y in comparison to indicated Ub species. For full spectra see **Appendix Fig. S4A**. **C)** Weighted chemical shift perturbation heat maps of phosphoUb L71Y in comparison to indicated Ub species. For full spectra see **Appendix Fig. S4B**. **D)** Lys11 resonance for Ub L71Y and phosphoUb L71Y in comparison to the split phosphoUb spectrum. **E)** Schematic of the Ub F4A mutation in which a residue from the neighbouring β1 strand modulates the Ub/Ub-CR equilibrium. **F)** Weighted chemical shift perturbation heat maps of Ub F4A in comparison to indicated Ub species. For full spectra see **Appendix Fig. S5A**. **G)** Weighted chemical shift perturbation heat maps of phosphoUb-CR F4A in comparison to indicated Ub species. For full spectra see **Appendix Fig. S5B**. **H)** Lys11 resonance for Ub F4A and phosphoUb F4A in comparison to the split phosphoUb spectrum.

Thus far the introduced mutations change residues on the slipping β5-strand. We wondered whether residues in the vicinity, e.g. from the neighbouring β1-strand, could also shift the observed equilibrium. A good candidate was Phe4 with its solvent exposed side chain (**Fig. 4E**), which would be anticipated to have only subtle effects on Ub conformation *per se*. Indeed, Ub F4A displayed a wild-type-like bTROSY spectrum (**Fig. 4H and F, Appendix Fig. S5A**). However, strikingly, phosphorylation of Ub F4A results in a spectrum where the most intense peaks are in a position associated with the phosphoUb-CR conformation and only a minor (∽12% by peak intensity) population of peaks match the common Ub conformation (**Fig. 4H and G, Appendix Fig. S5B**). This reverses the population ratio observed in the wt phosphoUb spectrum. This demonstrates that while the mutant resides in the common Ub conformation without phosphorylation, it almost completely shifts to a Ub-CR conformation upon phosphorylation. Hence, residues contacting and stabilising the slipping β5-strand are able to affect the conformational equilibrium.

**Comparative stability studies of Ub mutants**

The fascinating and unexpected conformational plasticity of Ub with regards to β5-strand slippage was further confirmed in comparative studies. We had previously shown decreased thermal stability of phosphoUb, which we speculated was due to the Ub/Ub-CR equilibrium (Wauer *et al*, 2015b). Indeed, differential scanning calorimetry (DSC) experiments revealed that Ub-CR mutants Ub L67S and Ub TVLN mutant display a *T*_*m*_ of ∽83 °C, with or without phosphorylation (**Fig. EV4)** (compared to 97 °C for wt Ub, and 87 °C for the phosphoUb equilibrium, (Wauer *et al*, 2015b)). In comparison, Ub F4A displays an intermediate stability (Tm 89 °C) consistent with NMR findings. Importantly, Ub L71Y is as stable as wt Ub (Tm 96 °C) indicating that the mutation does not induce unfolding, but merely stabilises the common Ub conformation. Hence, the Ub-CR conformation is less thermostable as compared to the common Ub conformation, and this explains lower stability of phosphoUb.

### Additional NMR evidence for a wt Ub-CR conformation

As already discussed above, the Ub-CR conformation results in a retracted C-terminus and an extended Ser65-containing loop, conferring new dynamic properties to the structure at these locations. As Ub mutants can modulate the newly found Ub / Ub-CR equilibrium, this prompted us to search for additional evidence for a Ub-CR conformation beyond CEST (**Fig. 1, Fig. EV1, Appendix Fig. S1**). For this, we used fast (ms timescale) hydrogen solvent exchange experiments known as clean chemical exchange (CLEANEX) and ^15^N{^1^H} heteronuclear NOE (hetNOE) experiments (that consider ps timescales of motion, **Fig. 1B**). CLEANEX experiments measure the ability of backbone amide protons to exchange with the solvent, thus reporting on the relative solvent exposure of each residue. CLEANEX experiments for each Ub variant revealed a similar set of solvent accessible residues for wt Ub, Ub L71Y and Ub F4A (**Appendix Fig. S6A and B)**, but considerably more solvent exchange especially in the Ser65-loop region in Ub TVLN. This is consistent with the structural data. Interestingly, residues of the nearby Leu8-loop report on the conformational preferences of each Ub mutant through their population averaged rates of solvent exchange (**Fig. 5A**). Note the loop in the TVLN mutant has the greatest degree of solvent accessibility, with the F4A mutant and wt Ub rates being greater than the L71Y Ub-CR inhibited mutant. This correlates with the overall stability seen in the *T*_*m*_ measurements (**Fig. EV4**).

**Figure 5.**
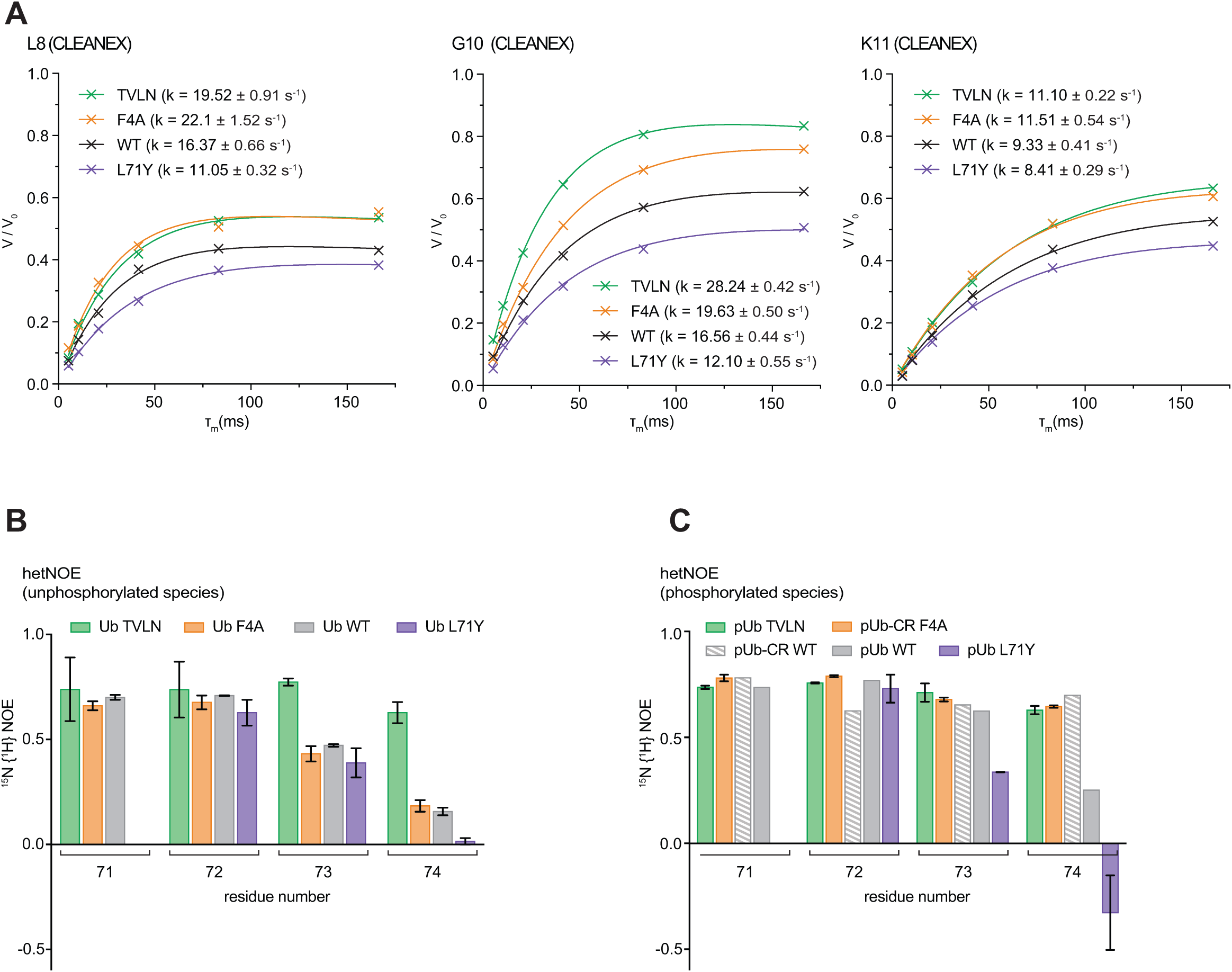
Comparative CLEANEX and hetNOE studies of Ub / Ub-CR mutants. **A)** CLEANEX experiments on Ub variants, comparing fitted rates of solvent exchange rates for selected residues of the Leu8-loop. See **Appendix Fig. S6A and B** for complete Ub CLEANEX rates and a graphical representation of the dataset. **B)** 15N {^1^H} hetNOE experiment for unphosphorylated Ub species, focussing at the flexible C-terminal tail. Large values are indicative of higher stability, and small or negative values indicate more flexibility. See **Appendix Fig. S7** for the complete datasets. **C)** As in B, for phosphorylated Ub variants. Data for pUb and pUb-CR WT is replotted from (Wauer *et al*, 2015b).

Further, indirect evidence for a contribution of the Ub-CR conformation in wt Ub was provided by hetNOE analysis, which assesses fast ps flexibility of individual residues (**Appendix Fig. S7A**). Probing the mobility of Arg74 with hetNOE experiments reveals an increased rigidity in Ub TVLN compared to wt Ub. In contrast the hetNOE values for Arg74 in the L71Y spectrum indicate increased flexibility as compared to the wt Ub spectrum (**Fig. 5B and C**). Given the above considerations one could infer that wt Ub has a contribution from its lowly populated Ub-CR conformation to the bulk dynamic properties. This would indicate that L71Y is the most ‘pure’ common Ub conformation of the mutants studied. While it is possible that Ub mutations contribute to some of the observed effects in CLEANEX and hetNOE analysis, the observed differences in stability and dynamics clearly also correlate with accessibility of the Ub-CR conformation.

### The Ub-CR conformation affects ubiquitination reactions

Our identification of Ub mutants adopting the Ub-CR conformation facilitated experiments to test the biochemical impact of this species on Ub assembly enzymes. As discussed above, retraction of the Ub C-terminal tail reduces flexibility of this key feature of Ub, but also disjoints the Ile44 hydrophobic patch, which is important for many enzymes of the assembly cascade.

Nonetheless, we found that Ub TVLN, which adopts the Ub-CR conformation in solution, was readily charged by E1 onto E2 enzymes, including UBE2D3, UBE2L3, UBE2S, UBE2N and UBE2R1 (**Fig. 6A**), which is perhaps surprising in light of recent findings that that the hydrophobic patch is important for E1-mediated E2 charging (Singh *et al*, 2017). However, the E1 reaction is known to be relatively permissive and can also accommodate conformation changing Ub mutations such as Ub L73P (Békés *et al*, 2013).

**Figure 6.**
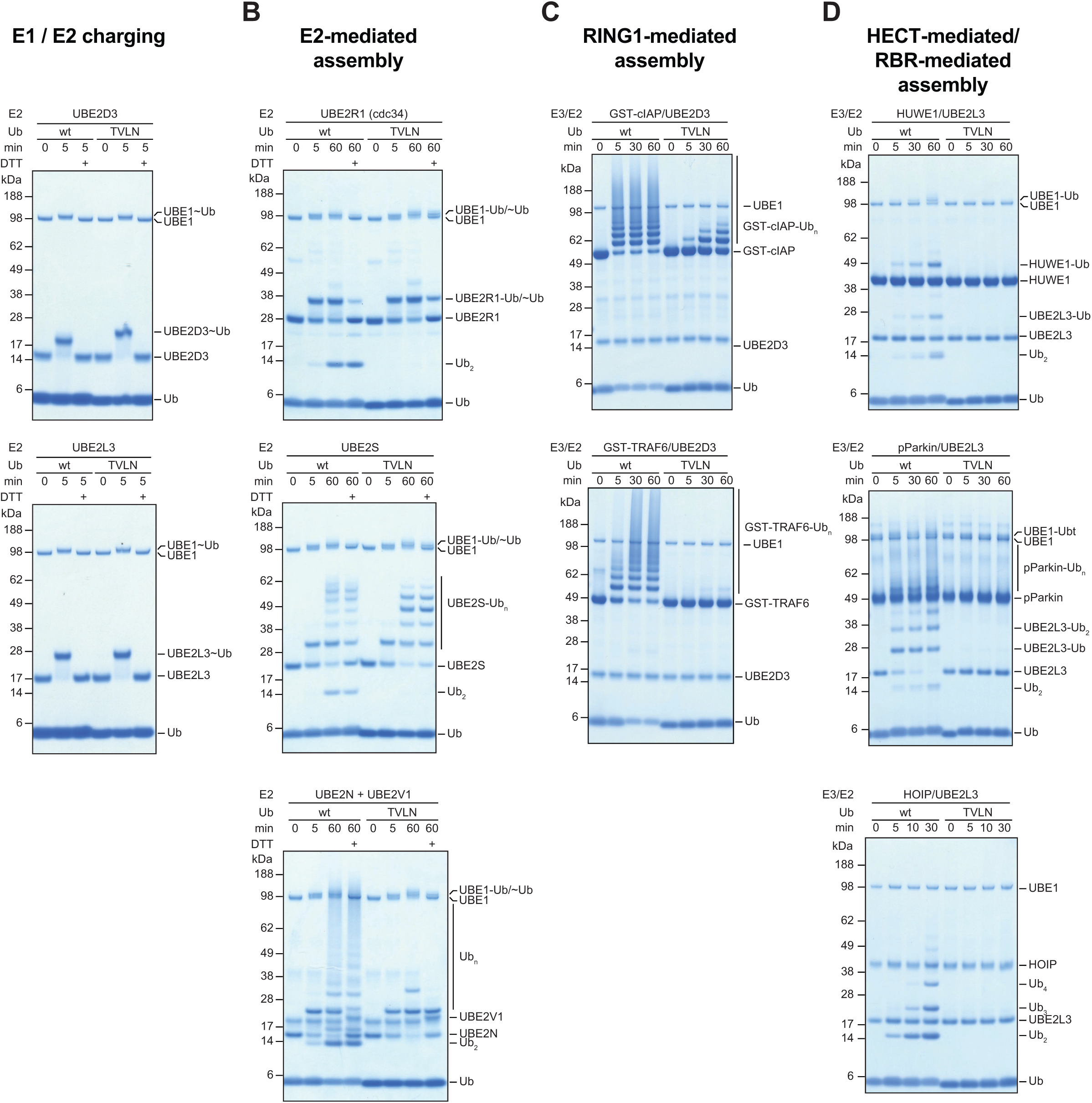
Effects of Ub-CR on ubiquitination reactions. Ubiquitination reactions were performed in parallel with wt Ub and the Ub-CR variant Ub TVLN at identical concentrations. Individual reactions were run for indicated times, resolved on 4-12% SDS PAGE gradient gels, and stained with Coomassie. **A)** E1 charging reactions with Ub and Ub TVLN on UBE2D3 (*top*) and UBE2L3 (*bottom*). **B)** E2-based Ub chain assembly reaction using UBE2R1 (*top*), UBE2S (*middle*) and UBE2N/UBE2V1 (*bottom*). E2 charging proceeds identically but chain assembly is inhibited with Ub TVLN, indicated by the lack of free diUb assembly. **C)** GST-E3-based autoubiquitination reaction with cIAP1 (aa 363-612) and TRAF6 (aa 50-211) in conjunction with UBE2D3. **D)** HECT- and RBR-based chain assembly in conjunction with UBE2L3. *Top*, HUWE1 HECT domain (aa 3993-4374). *Middle*, phosphorylated full-length Parkin. *Bottom*, RBR-LDD fragment of HOIP (aa 699-1072).

While charging was normal, Ub TVLN demonstrated impaired (UBE2S) or abrogated (UBE2R1, UBE2N/UBE2V1) E2-mediated chain assembly (**Fig. 6B**), and also impaired or abrogated chain assembly by RING E3 ligases (cIAP/UBE2D3, TRAF6/UBE2D3) (**Fig. 6C**), a HECT E3 ligase (HUWE1/UBE2L3), or RBR E3 ligases (Parkin/UBE2L3, HOIP/UBE2L3) (**Fig. 6D**). This shows that the Ub-CR conformation severely affects the Ub system. Consistently, in a large-scale mutational study in *Saccharomyces cerevisiae*, Ub L67S mutation was shown to have detrimental effects on yeast growth, despite being still charged onto E1 enzymes (Roscoe *et al*, 2013; Roscoe & Bolon, 2014). While Ub contains many essential residues and interfaces, our data suggest that the reported lack-of-fitness can be attributed to the Ub-CR conformation.

### The Ub-CR conformation stably binds *Ph*PINK1

While a Ub-CR-inducing mutation had the anticipated inhibitory effects on the ubiquitination cascade, we still wondered whether this conformation had physiological roles. A number of observations pointed towards potential importance in PINK1-mediated Ub phosphorylation. As discussed above, Ser65 in wt Ub is poorly accessible, but becomes more exposed in the Ub-CR conformation. Moreover, phosphorylation of Ub L67S and Ub TVLN mutants was markedly accelerated as compared to wt Ub as shown by qualitative Phos-tag gels (**Fig. 2E**). We hence tested how PINK1 interacted with its substrates, and performed bTROSY experiments with unlabelled *Ph*PINK1 (aa 115-575) and ^15^N-labelled wt Ub, Ub mutants, and Parkin Ubl domain (aa 1-76), respectively, in the absence of MgATP (**Fig. 7**). In the presence of *Ph*PINK1 all peaks were exchange broadened to some extent due to the formation of a weakly associated 62 kDa complex. A subset of Ub/Ubl peaks, which were additionally broadened, revealed the residues that interact with *Ph*PINK1 (**Fig. 7**, *left column*). The same samples were also used in CLEANEX experiments, whereby the binding of *Ph*PINK1 to Ub or Ubl masks the interacting residues on the substrate (**Fig. 7**, *right column*).

**Figure 7.**
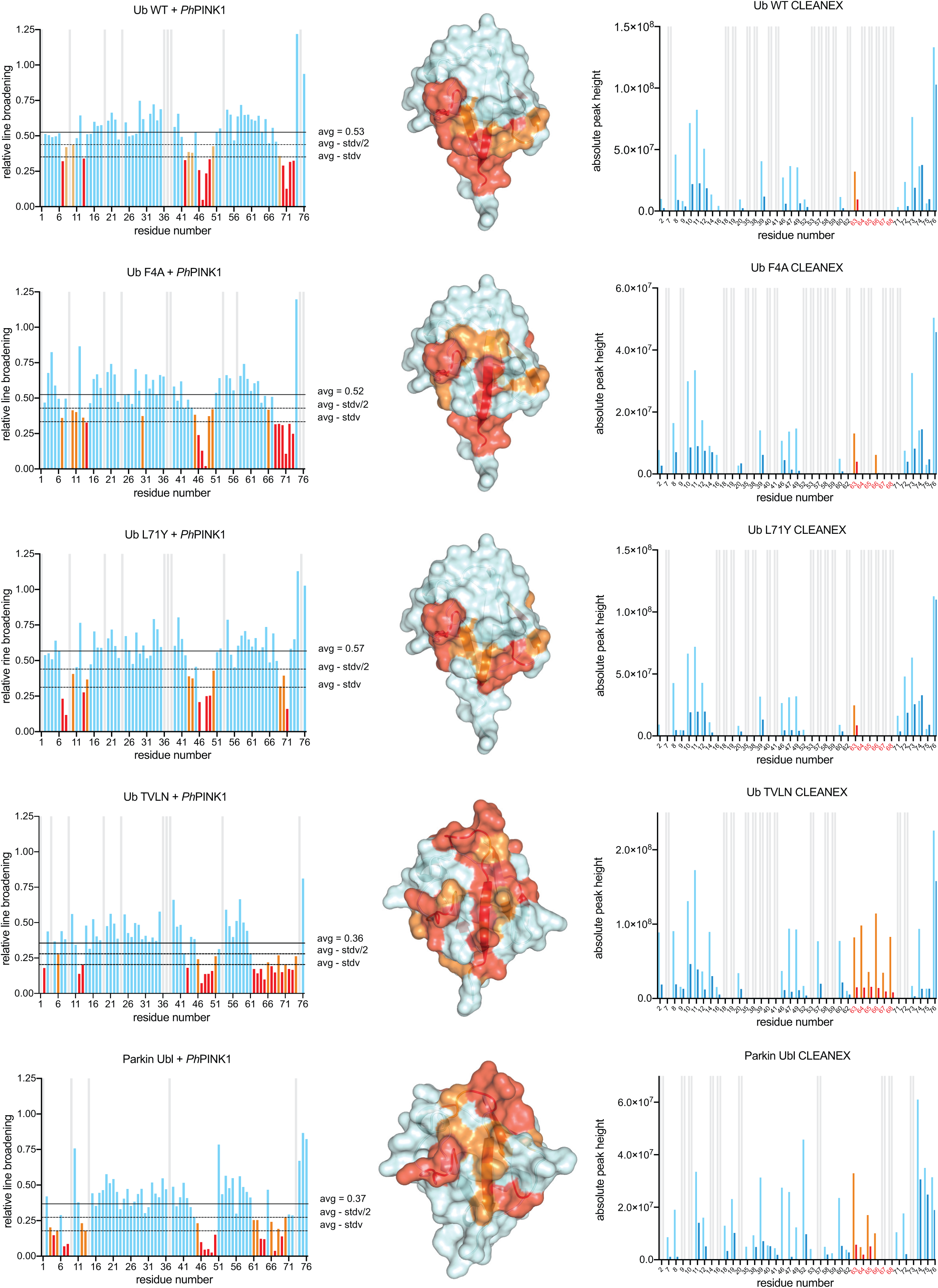
*Ph*PINK1 recognises the Ser65 loop in the Ub-CR conformation. (*Left column*) Complex formation between *Ph*PINK1 (without MgATP) and Ub variants or Parkin Ubl results in line broadening of NMR signals due to formation of a 62 kDa complex. Relative line broadening of Ub resonances is plotted with mean value indicated, and peak intensities decreasing by half or full standard deviation (stdv) from the (*Middle column*), Line broadened residues are plotted on the Ub surface. (*Right column*), CLEANEX experiments showing solvent exchanging residues (orange and light blue bars) on Ub variants and Parkin Ubl, and how these are affected by *Ph*PINK1 binding (red and blue bars). Orange/red bars highlight the Ser65-containing loop, which is exposed in Ub TVLN and highly protected after *Ph*PINK1 binding.

Remarkably, the footprint of *Ph*PINK1 on its substrates varied (**Fig. 7**, *middle column*). In wt Ub, as well as Ub F4A and Ub L71Y, broadened residues correspond to the C-terminal tail and the Ile44 patch, but strikingly did not include residues from the Ser65-containing loop. In these three samples, a similar degree of overall line broadening suggests similar (weak) binding. CLEANEX experiments of substrates without *Ph*PINK1 (light colours) and with *Ph*PINK1 (dark colours) reveal the Ile44 patch interaction of these substrates, but also confirm relative protection of the Ser65-loop, in which only Lys63 is sufficiently solvent exposed to be observed.

In contrast, the Ub TVLN mutant as well as the Parkin Ubl form larger interfaces involving the entire β5-strand, and importantly, all residues from the Ser65-containing loop. Moreover, overall line broadening was significantly stronger in Parkin Ubl and Ub TVLN samples as compared to wt Ub, suggesting that these substrates form a more stable complex. This was emphasised in the CLEANEX experiments collected for Ub TVLN, which in the apo state show the enhanced solvent accessibility of all resonances of the Ser65-loop. *Ph*PINK1 interaction leads to almost complete protection of the entire Ser65-loop of Ub TVLN showing that in the Ub-CR conformation the phosphorylation site can be part of the interface with PINK1.

Together, these experiments indicated that the significantly faster rate of Ub phosphorylation seen in the Ub-CR mutants such as Ub TVLN can be explained with enhanced binding of PINK1 to Ub-CR, which can form additional interactions via the Ser65-loop.

### Ub conformations affect PINK1 activity

The fact that Ub mutations stabilise the Ub-CR conformation in the absence of phosphorylation (**Figs. 2-5**), and the discovery that wt Ub dwells in a Ub / Ub-CR equilibrium under physiological conditions (**Fig. 1 and 5**), opened the fascinating possibility that the Ub-CR conformation is used or even required for PINK1-mediated Ub phosphorylation.

To test this, we compared phosphorylation rates by treating Ub, Ub mutants and Parkin Ubl samples with *Ph*PINK1 / MgATP, in qualitative experiments using Phos-tag gels (**Fig. EV5A)**, or in semi-quantitative, real-time experiments using ^15^N-labelled Ub/Ubl substrates in NMR experiments (**Fig. 8**). Direct assessment of individual peak disappearance/appearance over time from unphosphorylated to fully phosphorylated samples enabled generation of phosphorylation rate curves, revealing strikingly different rates (**Fig. 8**). The fastest rates were observed for Ub TVLN and Parkin Ubl, and these were almost indistinguishable from each other, even at lower enzyme concentrations (**Fig. EV5B**). 50% of the substrate was phosphorylated after ∽2 min. The Ub F4A sample was also quite fast, being half-phosphorylated after ∽5 min. In contrast, it took ∽90 min and ∽275 min to phosphorylate 50% of wt Ub or Ub L71Y, respectively, under identical conditions. Hence, phosphorylation of the Ub-CR mutant Ub TVLN is ∽45-140 fold faster as compared to variants where this conformation is much less populated (wt Ub) or disfavoured (Ub L71Y). This suggested that PINK1 not only prefers the Ub-CR conformation, but that it requires it for efficient phosphorylation of a Ub molecule.

**Figure 8.**
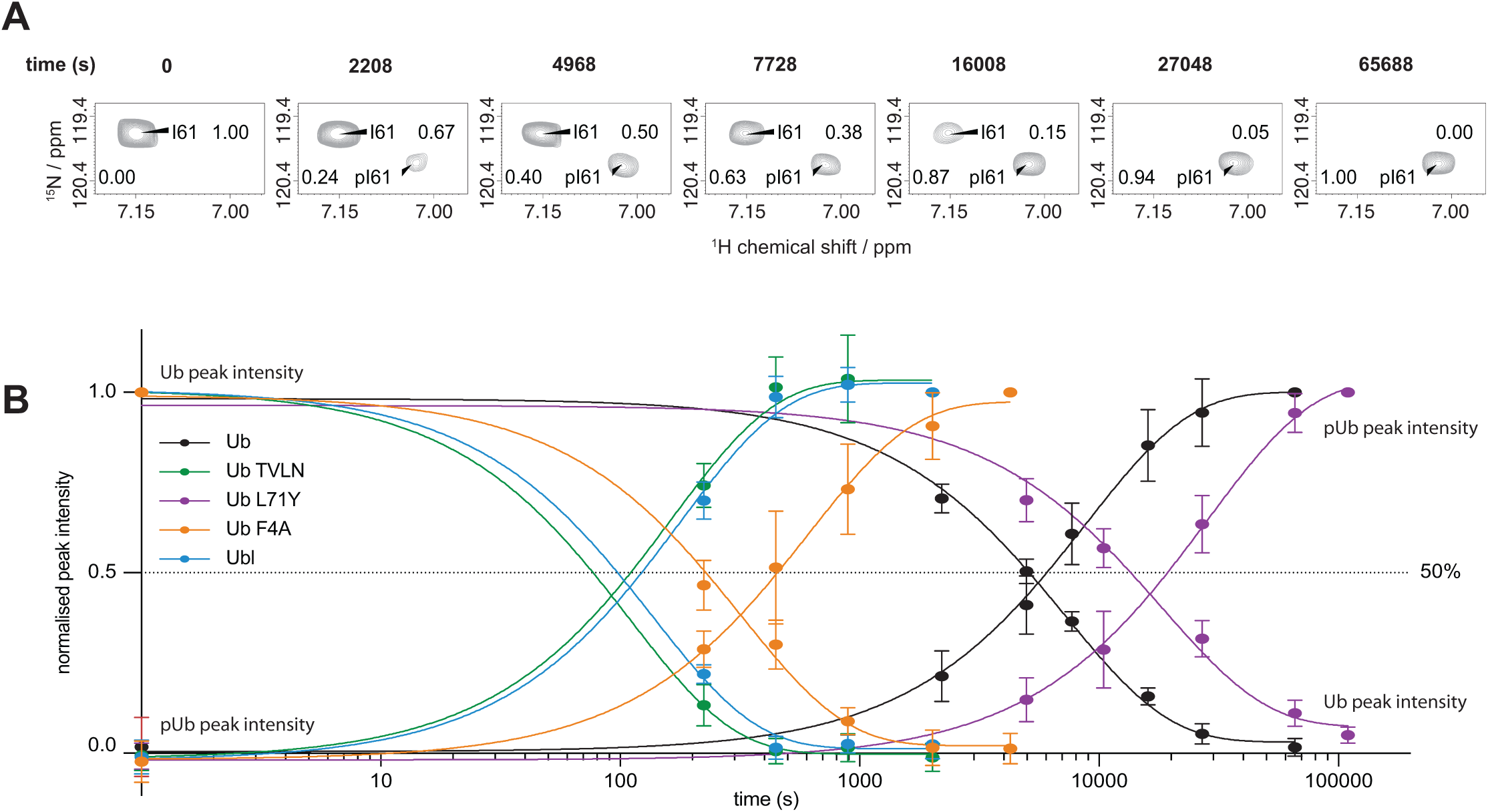
Ub in the Ub-CR conformation is a superior *Ph*PINK1 substrate. **A)** An *in-situ*, phosphorylation experiment was performed, in which suitable substrate signals were monitored for disappearance / appearance in an NMR time course as illustrated for Ile61. **B)** A series of bTROSY spectra were acquired in ∽4 (Ub TVLN, Ub F4A, and Ubl) or ∽8 min (wt Ub and Ub L71Y) increments following Ser65 phosphorylation by 350 nM *Ph*PINK1. Peak intensities of unphosphorylated and phosphorylated Ub/Ubl were normalised to the corresponding measurements in the initial (fully unphosphorylated) and final (fully phosphorylated) timepoints, respectively. Data from at least 9 individual resonances were averaged with error bars indicating standard deviation from the mean.

## DISCUSSION

Ubiquitin is a most fascinating molecule. Despite being the focus of three decades of biochemical, biophysical and structural research, we here uncover a new Ub conformation, in which the C-terminal β5-strand of Ub is retracted by two residues, which extends the upstream Ser65 containing loop, and shortens the otherwise extended Ub C-terminus. We show that Ub adopts the Ub-CR conformation under ambient conditions, and although this conformation is lowly populated, our data suggest that it is functionally relevant.

Our previous work showed that a very similar Ub-CR conformation was readily induced in Ser65-phosphoUb (Wauer *et al*, 2015b) which was recently confirmed by an NMR solution structure (Dong *et al*, 2017). We hence set out to to identify stable versions of phosphoUb-CR for further study. To our surprise, we identified point mutations that readily adopted the Ub-CR conformation even without phosphorylation, enabling us to shift the equilibrium. Still, we initially failed to detect a Ub-CR conformation in wt Ub, which was clearly a minute conformation not observable within the limits of datasets extending to μs dynamics of Ub (Lange *et al*, 2008). Based on the assumption that a potential wt Ub to Ub-CR exchange would occur at slower rates (as shown by the exchange rate of phosphoUb and phosphoUb-CR) and that the Ub-CR conformation would only be a lowly populated state referred to in the literature as a ‘dark’ or ‘invisible’ conformation (Baldwin & Kay, 2009; Kay, 2016), CEST experiments would be the analytical method of choice. Indeed, CEST experiments provided direct evidence for the existence of the Ub-CR conformation under physiologic conditions (phosphate buffered saline, at 37°C). This is an exciting finding that adds new complexity to the Ub conformational landscape.

Our mutational analysis explains previous findings that mutations of seemingly non-functional Ub residues severely affected Ub as well as cellular fitness (Roscoe *et al*, 2013; Roscoe & Bolon, 2014; Sloper-Mould *et al*, 2001). Most Ub mutations have to-date been explained with disruption of one of the various Ub binding interfaces (Komander & Rape, 2012). Whilst protein interactions are clearly a key function of Ub, we here reveal how some mutations may indirectly affect global Ub interaction capabilities by inducing a dysfunctional Ub-CR conformation.

Importantly, we also show a physiological role for a Ub-CR conformation. Ub is a well-folded, stable protein, and as such is an unlikely kinase substrate. Many protein kinases prefer or require disordered target sequences for phosphorylation. The well-ordered Ser65-containing loop in Ub does not fit this criterion, but the more mobile loop provided in the Ub-CR conformation enables efficient binding and phosphorylation. Hence, Ub-CR mutants are superior PINK1 substrates. Considering wt Ub, it is tempting to speculate that PINK1 stabilises the Ub-CR conformation, or indeed, that a Ub conformational change may impose a rate-limiting step for phosphorylation. So far, we have not observed this with wt Ub, but the timescales of binding experiments (μs) vs. conformational change (s) presents a challenge to directly detect a precatalytic state with wt Ub. It is exciting that we may be able to mimic this precatalytic state with the Ub TVLN mutant, and this may be useful for future structural studies on PINK1.

Our findings likely have pathophysiological relevance. PINK1 mutations result in AR-JP, and our results reveal that its key substrate, Ub, needs to be in a particular conformation to enable efficient phosphorylation. It is easy to imagine that conditions or binding partners that stabilise Ub in a common conformation (e.g. Ile44-patch binding domains in mitochondrial associated proteins) may impede PINK1 activity and imbalance the system. In this context, it will also be interesting to test whether different chain contexts modulate the observed Ub/Ub-CR equilibrium and affect the rate at which chains can be phosphorylated. A further open question relates to the Parkin Ubl domain, for which there is no evidence at current of a similar, C-terminally retracted Ubl conformation.

We had previously shown that the Ub-CR conformation is present also in phosphoUb (Wauer *et al*, 2015b), but strikingly, the only known phosphoUb receptor, Parkin, recognizes the common phosphoUb conformation and does not utilise the more distinctive phosphoUb-CR conformation (Wauer *et al*, 2015a). It is possible that alternative receptors for phosphoUb-CR exist, but it is also imaginable that this form exists predominantly to facilitate phosphorylation in the first place. While the Ub-CR conformation explains how Ser65 can be phosphorylated by PINK1, questions remain how other sites on Ub, such as well-ordered Thr12 and Thr14 on the β2-strand, can be phosphorylated.

More globally, our data explains how an inaccessible phosphorylation site in a folded protein can be targeted via exploitation of an ‘invisible’ conformation. Hence, our work is likely relevant for other kinases that target folded domains or proteins.

## METHODS

### Molecular biology

Ub constructs were cloned into pET17b vectors, and site directed mutagenesis was carried out using the QuikChange protocol with Phusion polymerase (NEB).UBE2D3, UBE2S, UBE2L3, UBE2R1, UBE2N/UBE2V1 full-length proteins and GST-cIAP1 (aa 363-612), GST-TRAF6 (aa 50-211) were expressed from pGEX6 vectors. Full-length *Hs*Parkin, *Ph*PINK1 (aa 115-575) and HOIP RBR-LDD (aa 699-1072) were expressed from a pOPIN-K vector, while *Hs*Parkin Ubl domain (aa 1-76) and HUWE1 catalytic domain (aa 3993-4374) were expressed from a pOPIN-S vector (Berrow *et al*, 2007). *Hs*Ube1/PET21d was a gift from Cynthia Wolberger (Addgene plasmid # 34965, (Berndsen & Wolberger, 2011)).

### Protein purification

All Ub mutants were expressed in Rosetta 2 (DE3) pLacI cells, and purified following the protocol of (Pickart & Raasi, 2005). In short, unlabelled proteins were expressed in 2xTY medium, protein expression was induced at OD_600_ of 0.6 - 0.8 with 200 μM IPTG and cells were harvested after 4 - 5 h at 37°C. Singly ^15^N labelled or doubly ^15^N and ^13^C labelled proteins were expressed in Minimal medium (M9 supplemented with 2 mM MgSO_4_, 50 μM ZnCl_2_, 10 μM CaCl_2_, trace elements, vitamins (BME vitamins solution, sterile filtered, Sigma)), supplemented with 1 g ^15^NH_4_Cl and 4 g glucose or ^13^C_6_ glucose where required. Protein expression was induced at OD_600_ of 0.5 - 0.6 with 200 μM IPTG and cells were harvested after O/N growth at 18°C.Labelled and unlabelled Parkin Ubl (aa 1-76) was expressed as a His-SUMO-fusion construct as described previously (Wauer *et al*, 2015b), and purified using HisPur™ Cobalt Resin (Thermo Fisher Scientific). The His-SUMO tag was cleaved using SENP1 during dialysis in cleavage buffer (25 mM Tris (pH 8.5), 300 mM NaCl, 2 mM β-mercaptoethanol) overnight at 4 °C. The His-SUMO tag was captured on HisPur™ Cobalt Resin. GST-tagged *Ph*PINK1 (aa 115-575), E2 and E3 enzymes were purified and Parkin phosphorylated as described earlier (Wauer *et al*, 2015b). For NMR studies the *Ph*PINK1 GST-tag was cleaved using Prescission protease. As a final step, all proteins subjected to NMR analysis were purified by SEC (Superdex 75 or Superdex 200, GE Life Science) in NMR-Buffer (18 mM Na_2_HPO_4_, 7mM NaH_2_PO_4_, 150 mM NaCl (pH 7.2) with 10mM DTT added for *Ph*PINK1 and Parkin Ubl). Proteins for biochemistry were purified by SEC (Superdex 75 or Superdex 200, GE Life Science) in 25 mM Tris, 150 mM NaCl (pH 7.4)).

### Phos-tag assays

Phosphorylation of Ub constructs and Parkin Ubl was performed by incubating 15 μM substrate with indicated GST-*Ph*PINK1 concentrations in 25 mM Tris (pH 7.4), 150 mM NaCl, 10 mM MgCl_2_, 10 mM ATP, 1 mM DTT at 22 °C or 37 °C as indicated.Reactions were quenched at the given time points with EDTA-free LDS sample buffer.

Samples were analysed by Mn^2+^ Phos-tag SDS-PAGE. A 17.5% (w/v) Acrylamide gel was supplemented with 50 μM Phos-tag AAL solution (Wako Chemicals) and 50 μM MnCl_2,_ and stained with Instant Blue SafeStain (Expedeon). An EDTA-free Tris-Glycine running buffer was used.

### Mass-spectrometry analysis

LC-MS analysis was carried out on an Agilent 1200 Series chromatography system coupled to an Agilent 6130 Quadrupole mass spectrometer. Samples were eluted from a phenomenex Jupiter column (5 μm, 300 Å, C4 column, 150 x 2.0 mm) using an acetonitrile gradient + 0.2% (v/v) formic acid. Protein was ionized using an ESI source (3 kV ionization voltage) and spectra were analysed in positive ion mode with a mass range between 400-2,000 m/z. Averaged spectra were deconvoluted using the manufacturer’s software and plotted using GraphPad Prism (version 7).

### Ub phosphorylation by *Ph*PINK1

Purified Ub variants were incubated at a 100:1 ratio with *Ph*PINK1 in phosphorylation buffer (10 mM ATP, 20 mM Tris (pH 7.4), 10 mM MgCl_4_, 150 mM NaCl, 1 mM DTT). Reaction progress at 25°C was monitored using LC-MS, and once there were no changes in recorded spectra, the reaction mixture was dialysed against water, using a 3.5 kDa cut-off Dialysis cassette (Thermo scientific). The dialysate was applied to an anion exchange (MonoQ 5/50 GL, GE Life Sciences) column. PhosphoUb was eluted by 50 mM Tris (pH 7.4) and further purified by SEC (Superdex 75, GE Life Sciences) into NMR-buffer. PhosphoUb TVLN for crystallography was purified by SEC in 25 mM Tris (pH 7.4).

### Crystallisation, data collection and structure determination

Ub L67S was crystallised at 12.5 mg/ml by sitting-drop vapour diffusion against 3 M (NH_4_)_2_SO_4_, 0.1 M MES (pH 6.0) using a 2:1 protein to reservoir ratio at 18°C. A single crystal was harvested and vitrified in liquid nitrogen.

PhosphoUb TVLN was crystallised at 11.2 mg/ml by sitting-drop vapour diffusion against 3.2 M (NH_4_)_2_SO_4_, 0.1 M bicine (pH 9.0), in a 1:1 protein to reservoir ratio at 18°C. A single crystal was harvested and vitrified in liquid nitrogen. Ub L67S diffraction data was collected at the Diamond Light Source, beam line I-04, while phosphoUb TVLN was collected on an FR-E^+^ SuperBright ultra high intensity microfocus rotating copper anode (λ=1.5418A°) generator equipped with a MAR345 detector. Diffraction data were processed with iMosflm (Battye *et al*, 2011) and scaled with AIMLESS (Evans, 2006). Structures were determined by molecular replacement, using wt Ub (pdb-1UBQ, (Vijay-Kumar *et al*, 1987)) aa 1-59 as a search model in Phaser (McCoy *et al*, 2007). Iterative rounds of model building and refinement were performed with Coot (Emsley *et al*, 2010) and PHENIX (Adams *et al*, 2011), respectively. All structural figures were generated in Pymol (www.pymol.org).

Data collection and refinement statistics can be found in **Table 1**.

### Stability measurements

Samples were dialysed into NMR buffer (18 mM Na_2_HPO_4_, 7 mM NaH_2_PO_4_, 100 mM NaCl (pH = 7.2) using 3.5 kDa MW cut off dialysis cassettes (Thermo Scientific) and subsequently diluted to 50 μM. Differential scanning calorimetry (DSC) was performed using a VP-Capillary DSC instrument (Malvern Instruments). Samples were scanned at a heating rate of 90°C / h in mid-feedback mode. Data were corrected for instrumental baseline using average buffer scans recorded immediately before and after Ub runs and plotted. After concentration normalisation, the intrinsic protein baseline between pre- and post-transitional levels was corrected using the progress function in the Origin software supplied with the instrument. Corrected endotherms were fitted to a non-two state model allowing *T*_*m*_, *ΔH* calorimetric and *ΔH* van’t Hoff to vary independently.

### Ubiquitination assays

Ubiquitination assays were essentially performed according to (Wauer *et al*, 2015b), with reactions performed in ubiquitination buffer (30 mM HEPES (pH 7.5), 100 mM NaCl, 10 mM ATP, 10 mM MgCl_2_). For E2 charging and E2-mediated assembly *Hs*UBE1 was used at 0.2 μM, Ub was used at 20 μM and E2s were used a 4 μM. For E3-mediated assembly *Hs*UBE1 was used at 0.2 μM, Ub was used at 20 μM, E2s were used a 2 μM and GST-cIAP1, GST-TRAF6, HUWE1, pParkin were used at 5 μM, while HOIP RBR LDD was used at 1 μM. Samples were taken at indicated time points, the reactions quenched with LDS sample buffer with reducing agent unless otherwise indicated, resolved on 4-12% SDS Gradient gels (NuPage) and stained with Instant Blue SafeStain. A representative example of an experiment done at least in duplicate is shown.

## NMR

### General acquisition parameters

NMR acquisition was carried out at 25 °C unless otherwise stated on either Bruker Avance III 600 MHz, Bruker Avance II+ 700 MHz or Bruker Avance III HD 800 MHz spectrometers equipped with a cryogenic triple resonance TCI probes. Topspin (Bruker) and NMRpipe (Delaglio *et al*, 1995) were used for data processing and Sparky (T. D. Goddard and D. G. Kneller, SPARKY 3, UCSF, http://www.cgl.ucsf.edu/home/sparky/) was used for data analysis. ^1^H, ^15^N 2D BEST-TROSY experiments (band selective excitation short transients transverse relaxation optimised spectroscopy) were acquired with in-house optimised Bruker pulse sequences incorporating a recycling delay of 400 ms and 1024*64 complex points in the ^1^H, ^15^N dimension, respectively. High quality data sets were collected in approximately 9 min.

### Backbone chemical shift assignments

*De novo assignments or reassignments (L67S, TVLN, pTVLN, F4A, pF4A)* NMR acquisition was carried out at 25 °C on Bruker Avance III 600 MHz spectrometer equipped with a cryogenic triple resonance TCI probe. Backbone chemical shift assignments were completed using Bruker triple resonance pulse sequences. HNCACB spectra were collected with 512*32*55 complex points in the ^1^H, ^15^N,^13^C dimensions respectively. CBCA(CO)NH, HNCO and HN(CA)CO spectra were collected with 512*32*48 complex points in the ^1^H, ^15^N, ^13^C dimensions respectively. All experiments were collected using Non Uniform Sampling (NUS) at a rate of 50% of complex points in the ^1^H, ^15^N, ^13^C dimensions respectively and reconstructed using compressed sensing (Kazimierczuk & Orekhov, 2011). Assignment of the common conformation peaks seen in the pF4A ^15^N-^1^H spectra was aided by analysis of ZZ-exchange experiments (Latham *et al*, 2009) collected with 50, 75, 150, 200, 400 and 800 ms delays using the Bruker 950 MHz Avance III HD spectrometer at the MRC Biomedical NMR centre for optimised sensitivity.Due to the similarity of the L71Y and pL71Y HSQC spectra to WT-Ub cross peak assignment was simply confirmed by analysis of a ^15^N NOESY-HSQC collected with a mixing time of 120 ms and 1024*32*48 complex points in the ^1^H, ^15^N and ^1^H dimensions.

Previously published assignments of peaks in the parkin (1-76) Ubl and pUbl by (Aguirre *et al*, 2017) were downloaded from the BioMagResBank www.bmrb.wisc.edu (accession number 30197). Weighted chemical shift perturbation calculations were performed using the following relationship: ((Δ^1^H)^2^+(Δ^15^N/5)^2^)^0.5^ where the Δ denotes the difference in ppm of the chemical shift between the peaks of phosphorylated and non phosphorylated peaks of the same ubiquitin or between different ubiquitin species. Data were plotted with GraphPad Prism (version 7).

### ^15^N {^1^H }-heteronuclear NOE measurements

^15^N {^1^H }-heteronuclear NOE (hetNOE) measurements were carried out using standard Bruker pulse programs, applying a 120° ^1^H pulse train with a 5 ms inter-pulse delay for a total of 5 s interleaved on- or off-resonance saturation. The hetNOE values were calculated from peak intensities according the equation I_on_/I_off_.

### CLEANEX experiment on Ub (Fitting)

All CLEANEX experiments were collected at 800 MHz with a 3 s acquisition delay and mixing times of 5.2, 10.4, 20.8, 41.6, 83.2 and 166.4 ms using standard Bruker pulse programs. Backbone amide protons that exchanged with the bulk solvent were fitted using established methods (Hwang *et al*, 1998), with exchange rates plotted using GraphPad Prism (version 7).

## CEST

^15^N Pseudo-3D CEST experiments were collected at 700 MHz at 25, 37 and 45°C using established pulse sequences (Vallurupalli *et al*, 2012). At each temperature experiments were acquired with an exchange period of 400 ms and a weak B_1_ spinlock field of either 12.5 or 25 Hz, which was calibrated according to (Vallurupalli *et al*, 2012) and applied in a range between 102-134 ppm at 184 or 92 frequency points respectively. ^15^N CEST profiles were plotted as I/I_0_ against applied B1 field, with the I_0_ value taken as first slice where the exchange period was omitted.

### PhPINK1 – Ub/Ubl binding experiments

Binding experiments were performed by recording BEST-TROSY and CLEANEX (with 166.4 ms mixing time) with 65 μM Ub/Parkin Ubl constructs with and without equimolar amounts of *Ph*PINK1. For the BEST-TROSY experiments, the peak heights of the datasets with *Ph*PINK1 were normalised against the respective peaks without *Ph*PINK1 for each Ub / Parkin Ubl construct and plotted accordingly. For the CLEANEX experiments, the absolute peak heights with and without *Ph*PINK1 were plotted side by side.

### Phosphorylation rate measurements by NMR

Phosphorylation was performed by incubating 100 μM labelled substrate (Ub or Parkin Ubl) with 350 nM *Ph*PINK1 in NMR-Buffer supplemented with 10 mM MgCl_2_/ATP at 25 °C. 700 MHz BEST-TROSY experiments were carried out to monitor phosphorylation with 8 scans and 128 increments for wt Ub and Ub L71Y (∽8 min), and 4 scans and 100 increments for Ub F4A, Ub TVLN and Parkin Ubl (∽3.5 min). To compare Ub TVLN and Parkin Ubl phosphorylation rates, 65 μM Ub TVLN or Parkin Ubl were incubated with 20 nM *Ph*PINK1 in NMR-Buffer and 10 mM MgCl_2_/ATP at 25 °C. 600 MHz BEST-TROSY experiments were recorded with 8 scans and 128 increments. Peak heights of each time point were normalised against the peak height of the first (no phosphorylation) and last (full phosphorylation) time point, respectively. A minimum set of 9 peaks for each construct was used to plot phosphorylation rates (wt Ub: I3, F4, I13, T14, L15, E18, I23, V26, K29, I30, L43, I44, F45, G47, L50, E51, D52, S57, N60, I61, K63, E64, L67, H68; Ub TVLN: Q2, K6, T7, L15, I61, K63, E64, V66, N67, H68; Parkin Ubl: F4, R6, E16, S22, C59, D60, Q64, H68, V70; Ub F4A: K27, A28, K29, I30, D32, Q41, K48, L50, D52, L71; Ub L71Y: Q2, V5, K6, T14, L15, I23, K29, I44, F45, G47, L50, L56, S57, N60, I61, Q62, E64, S65, T66, L67, H68, V70, R72, L73) and a set of 4 peaks for the Ub TVLN and Parkin Ubl phosphorylation rate comparison (Ub TVLN: I3, K6, T7, Q62; Parkin Ubl: E16, Q25, K27, E28, F45, K48, E49, D60, Q64, V67).

Data were plotted with GraphPad Prism (version 7).

## Acknowledgements

We would like to thank Minmin Yu and beam-line staff at Diamond Light Source, beam lines I-04. Access to DLS was supported in part by the EU FP7 infrastructure grant BIOSTRUCT-X (contract no. 283570). We further thank Tom Frenkel and facility staff at MRC Biomedical NMR Centre for 950 MHz NMR data collection. This work was supported by the Francis Crick Institute through provision of access to the MRC Biomedical NMR Centre. The Francis Crick Institute receives its core funding from Cancer Research UK (FC001029), the UK Medical Research Council (FC001029), and the Wellcome Trust (FC001029). We also thank Chris Johnson and the LMB Biophysics facility for their assistance with the DSC measurements. We thank members of the DK lab for reagents and discussions. This work was supported by the Medical Research Council [U105192732], the European Research Council [309756], and the Lister Institute for Preventive Medicine (to DK).

## Accession numbers

Coordinates and structure factors for have been deposited with the protein data bank accession codes 5OXI (Ub L67S), 5OXH (phosphoUb TLVN).

## Author contribution

Conceptualisation and experimental design, CG, AS, JLW, SMVF, DK; investigation, CG, AS, JLW, CMJ, JNP; writing, DK; funding acquisition, DK. All authors commented on and improved the text.

---

**Extended View Figure 1.**
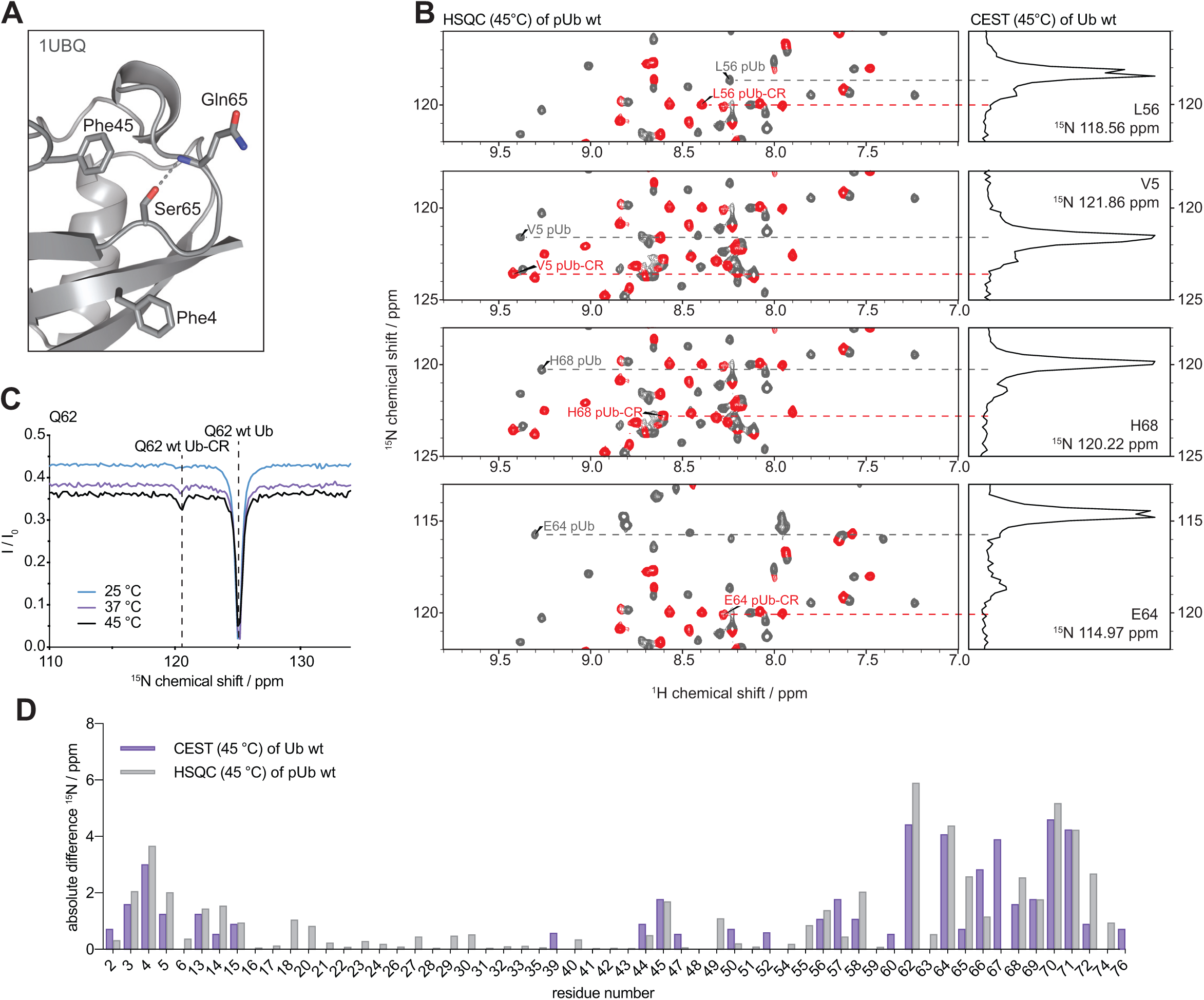
CEST characterisation of wt Ub. **A)** Structure of the Ser65 loop in the common conformation of wt Ub (1UBQ, (Vijay-Kumar *et al*, 1987)), also showing nearby aromatic residues. The Ser65 hydrogen bond is indicated. **B)** Additional CEST peaks as shown in **Fig. 1C**. For full spectra see **Appendix Fig. S1**. **C)** Temperature dependence of CEST experiments for a selected resonance (Gln62). A Ub-CR CEST peak is clearly observed at 37 °C but more pronounced at 45 °C. **D)** Absolute ^15^N-positional difference plot for all resonances according to Ub CEST at 45 °C, in direct comparison to the ^15^N-positional difference for each phosphoUb / phosphoUb-CR resonance pair in the phosphoUb HSQC spectrum at the same temperature. Extended View Figure EV2

**Extended View Figure 2.**
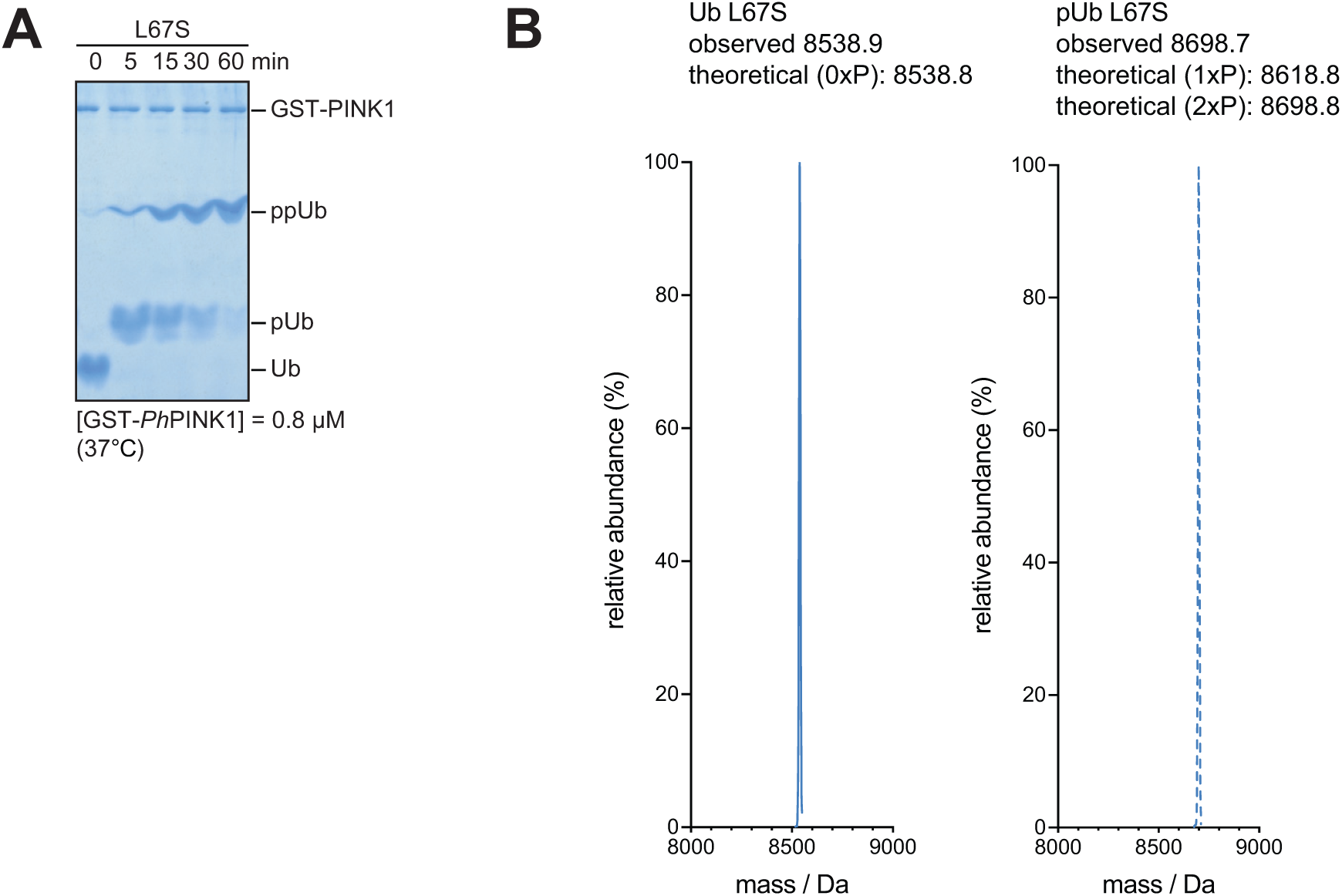
Phosphorylation of Ub L67S. **A)** Phos-tag gel following Ub L67S phosphorylation by *Ph*PINK1. Ub L67S is phosphorylated twice. **B)** LC-MS intact mass analysis of Ub L67S in unphosphorylated (left) and phosphorylated state (right). Two phosphorylation sites are observed.

**Extended View Figure 3.**
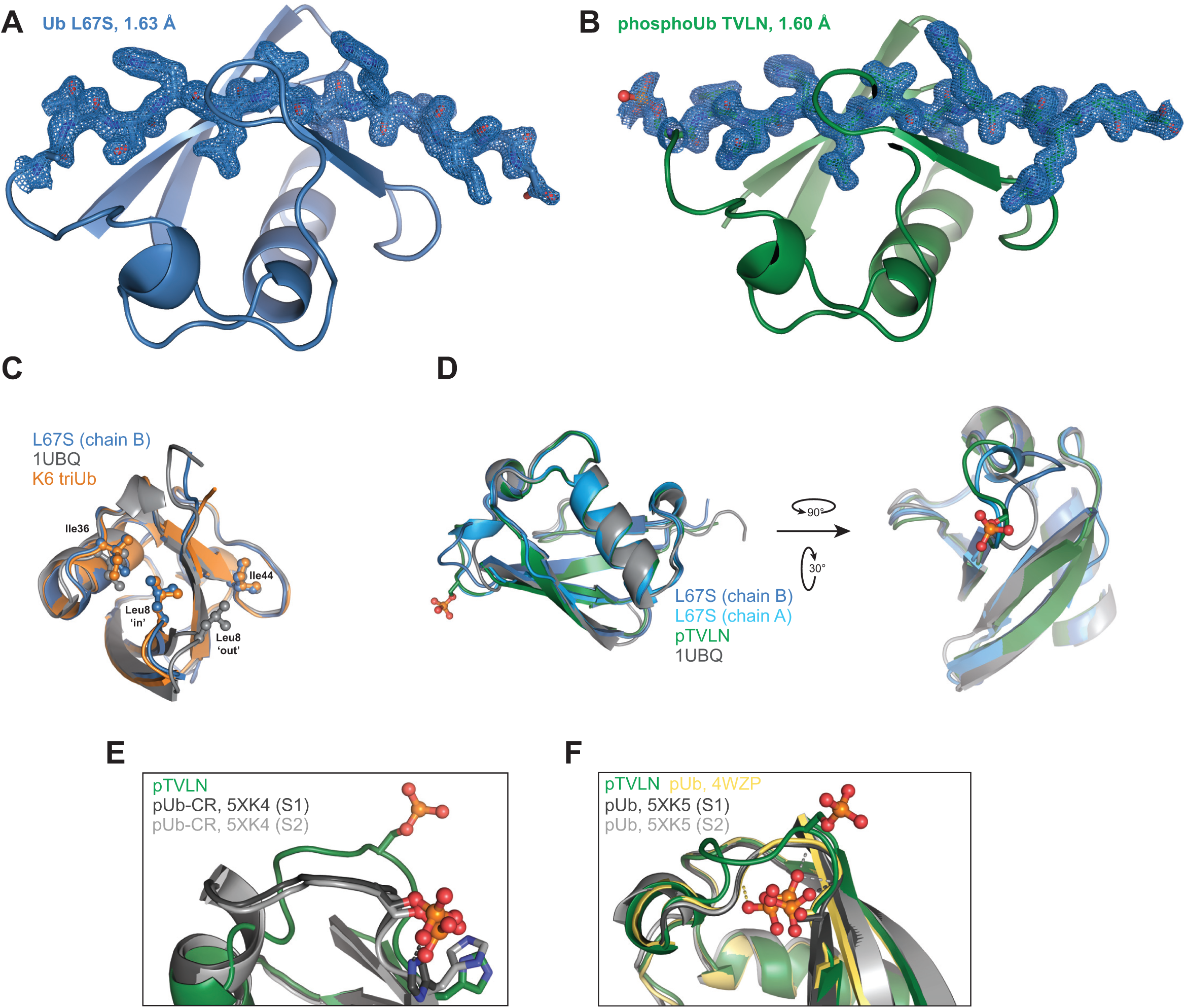
Ub-CR crystal structures and comparison. **A)** Ub L67S in a view as in **Fig. 3A**, showing 2|F_o_|-|F_c_| electron density at 1σ for the β5-strand. **B)** As in A, for phosphoUb TVLN. **C)** Superposition of Ub with distinct positions of the β1-β2 loop. Unlike the common Ub conformation, where this loop is predominantly in the ‘out’ position and Leu8 contributes to the Ile44 patch, the β1-β2 loop is in the ‘in’ position in Ub-CR mutants such as Ub L67S; in this position Leu8 contributes to the Ile36 patch. This loop conformation has also been observed in a subset of crystal structures, such as in Lys6-linked triUb (PDB-id 3zlz). See (Hospenthal *et al*, 2013) for further analysis. **D)** Two additional views of Ub-CR structures, highlighting the differences in the Ser65-containing loop. **E)** Comparison of phosphoUb TVLN with NMR structures of phosphoUb-CR (PDB-id 5xk4, (Dong *et al*, 2017)). **F)** Comparison of phosphoUb TVLN and phosphoUb crystal structures (pdb-id 4wzp, (Wauer *et al*, 2015b) with NMR structures of phosphoUb (pdb-id 5xk5, (Dong *et al*, 2017)).

**Extended View Fig. 4.**
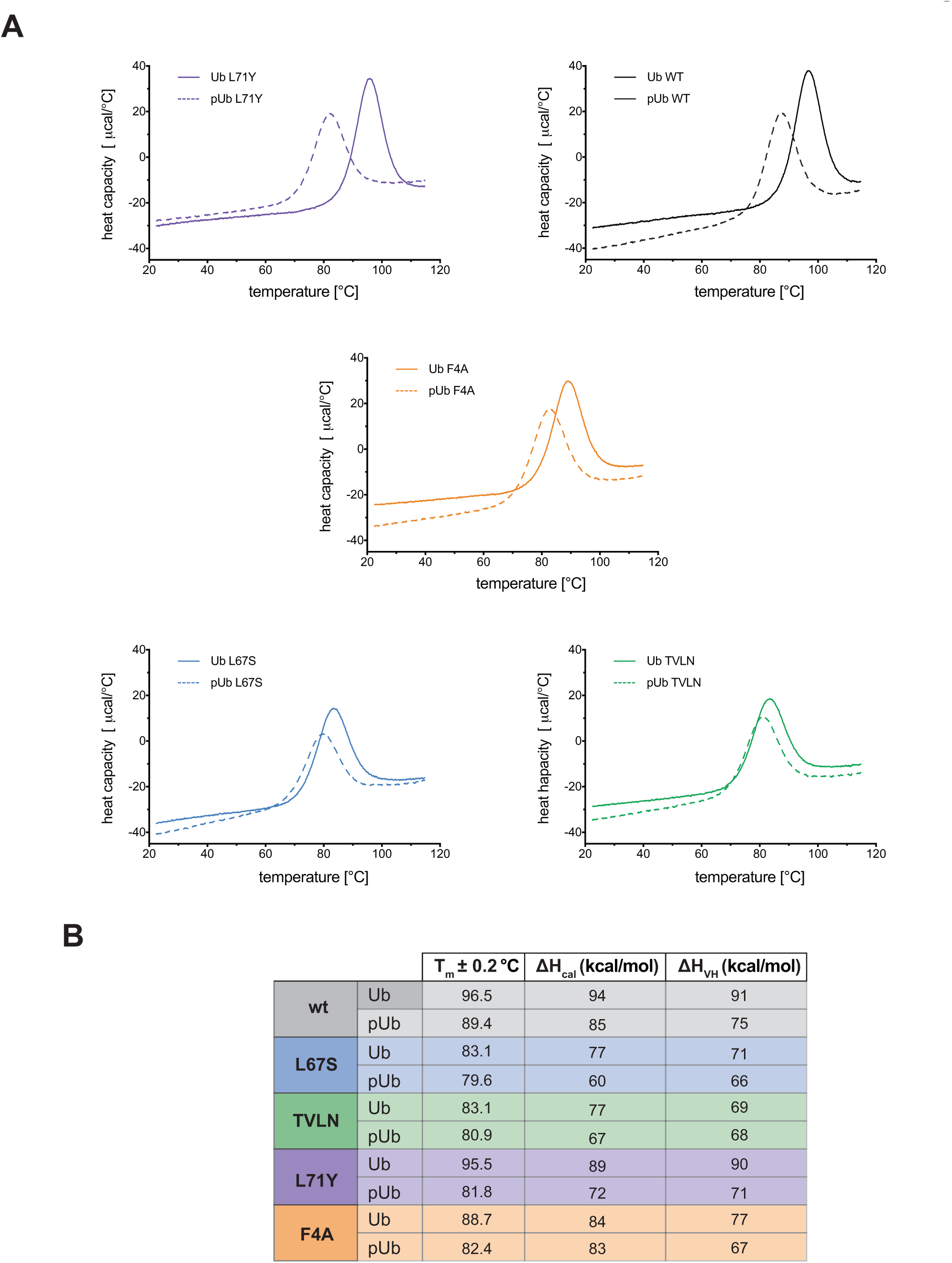
Stability measurement of Ub-CR variants. **A)** Differential scanning calorimetry endotherms of indicated Ub variants in unphosphorylated and phosphorylated form. **B)** Table summarising fitted melting parameters of each species. Similar values for ΔH_cal_ and ΔH_van’t Hoff_ indicate lack of intermediates during unfolding.

**Extended View Figure 5.**
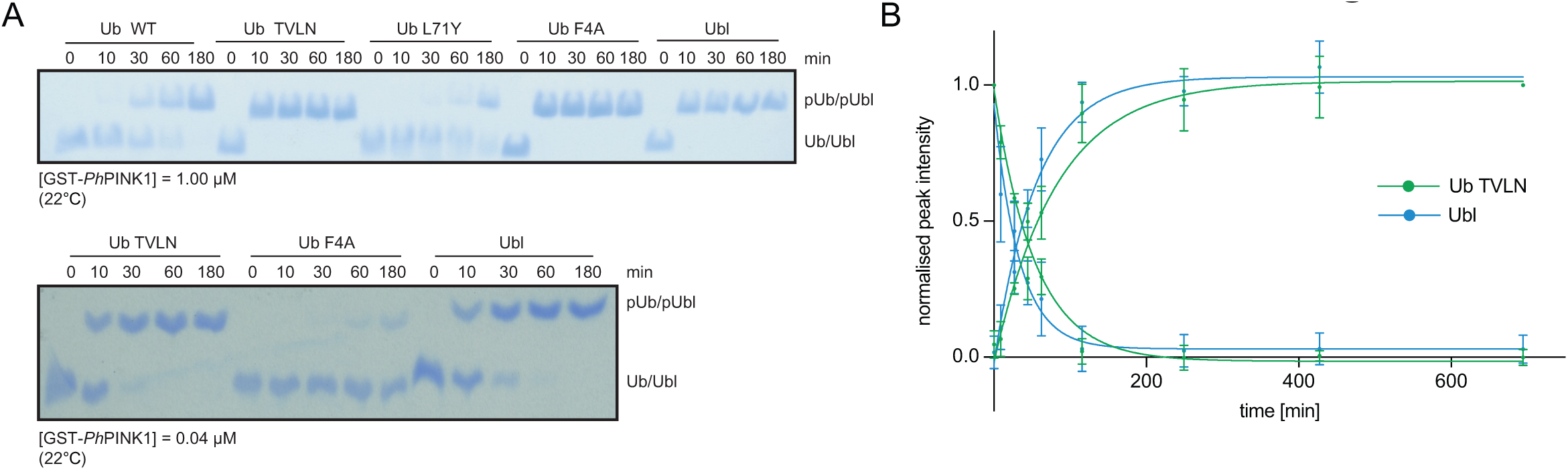
Ub phosphorylation experiments by Phos-tag analysis. **A)** Analysis of substrate phosphorylation by *Ph*PINK1, using Phos-tag gels. A lowed concentration of *Ph*PINK1 is used to reveal similar kinetics of Ub TVLN and Parkin Ubl phosphorylation (*bottom gel*). **B)** NMR-based Ub TVLN and Parkin Ubl phosphorylation as in **Fig. 8**, but at lower *Ph*PINK1 enzyme concentration of 20 nM.

